# Rapid pronucleus assembly using cytoplasmic RNAs in fertilized eggs of *Xenopus laevis*

**DOI:** 10.1101/2025.02.19.636958

**Authors:** Mizuki Ikeda, Yuto Tanaka, Tatsuya Shohoji, Yuki Hara

## Abstract

The size of the nucleus, which serves as the site for essential cellular functions such as replication and transcription, is dynamically altered to support these functions in response to the surrounding environment. During the brief cleavage period in metazoan embryos, the small, hypercondensed sperm nucleus with silenced chromatin undergoes a dramatic transformation into a large, round pronucleus with relaxed chromatin, enabling the activation of chromatin functions necessary for subsequent development. However, it remains unclear whether the egg cytoplasm-specific molecular environment plays a role in pronucleus assembly. In this study, we evaluated the impact of abundant RNAs in eggs on pronucleus assembly by utilizing a cell-free reconstruction of interphase nuclei in *Xenopus laevis* egg extract. We found that when RNA levels deviated from the conventional concentration, the growth rate of the interphase nucleus decreased. Additionally, the addition of RNAs led to a more dispersed chromatin distribution and the dissociation of sperm-specific nuclear proteins from the chromatin. These chromatin remodeling properties, which were reproducible with the introduction of cationic compounds, facilitated the incorporation of somatic histones into the chromatin in reconstructed nuclei. Based on these findings, we propose that cytoplasmic RNAs promote the rapid decondensation of negatively charged chromatin from a hypercompacted state and the removal of positively charged protamines from sperm chromatin via electrical interactions. This remodeling accelerates pronucleus assembly during the brief cleavage period following fertilization and promotes the rapid growth in nucleus size.

## Introduction

In eukaryotic cells, the nucleus compartmentalizes genomic DNA using lipid bilayer membranes that separate it from the cytoplasm. The size of the nucleus is generally altered in response to the cellular environment, such as during cell cycle progression, developmental processes, and evolutionary processes (Jorgensen *et al*., 2007; Neumann and Nurse, 2007; Hara, 2020; Wesley *et al*., 2020; Malerba and Marshall, 2021). For instance, nuclear size increases with cell size during interphase, maintaining a positive correlation between nuclear size and cell size. The regulation of nuclear size is involved in controlling genetic functions, such as transcription and chromosome condensation (Hara *et al*., 2013; Jevtić and Levy, 2015), as well as non-genetic functions, including cell migration and mechanosensing (Lomakin *et al*., 2020; Tollis *et al*., 2022). Abnormal nuclear size, which is pathologically observed in cancer cells, correlates with malignancy and metastasis in some tumor cells (de Andrea *et al*., 2011), highlighting the physiological importance of nuclear size control. Therefore, the size of the nucleus must be controlled to adapt its nuclear functions to the cellular environment.

Dynamic control of nuclear size is essential in the embryos of metazoan species after fertilization. During this period, highly compacted sperm chromatin with silenced chromatin functions, such as replication and transcription, is converted into a nucleus with relaxed chromatin and active chromatin functions within a relatively short interphase. For example, in the clawed frog *Xenopus laevis*, the volume of the sperm nucleus increases by approximately 1,000-fold to form the male pronucleus, which exceeds 10,000 µm^3^ within 1.5 h after fertilization. At the same time, a sperm-specific nuclear protein called protamine, which is key to chromatin packing, is converted into a conventional histone-containing nucleosome. Two molecular mechanisms that regulate these drastic structural changes in the nucleus and chromatin after fertilization have been identified. The first mechanism is mediated by the chaperone protein nucleoplasmin. Nucleoplasmin in the egg cytoplasm facilitates the exchange of sperm-specific protamines for somatic histones in chromatin (Ohsumi and Katagiri, 1991; Philpott *et al*., 1991; Philpott and Leno, 1992). The second mechanism accelerates the growth rate of the interphase nucleus. In fertilized eggs, the supply of nuclear constituents from the cytoplasm increases.

Lamins, which are the main constituent proteins of the nuclear lamina underneath the nuclear envelope in vertebrates (Levy and Heald, 2010; Jevtić *et al*., 2015; Brownlee and Heald, 2019), and lipid membranes, which compose the nuclear membrane (Hara and Merten, 2015; Kume *et al*., 2019; Mukherjee *et al*., 2020), have been identified as determinant constituents. Lamins and lipid membranes are imported from the cytoplasm via the nuclear import system (Levy and Heald, 2010) and transported toward the nucleus along microtubules (MTs) by motor proteins such as dyneins (Wang *et al*., 2013), respectively. The amount of these determinants supplied to the nucleus positively correlates with cell size (Levy and Heald, 2010; Hara and Merten, 2015; Brownlee and Heald, 2019; Mukherjee *et al*., 2020), so the fertilized egg, being the largest during embryonic cleavage, can maximize the growth rate of the nucleus. In addition to the supply of constituents, the chromatin structure within the nucleus is also known to play a role in nuclear size control. Manipulating the condensation state of chromatin within the nucleus by adding drugs to alter histone modifications or induce hypercondensation coincides with changes in nuclear growth rate and nuclear stiffness, which occur through the alteration of the repulsive force of negatively charged chromatin (Shimamoto *et al*., 2017; Stephens *et al*., 2019; Heijo *et al*., 2020; Schibler *et al*., 2023).

The molecular environment and physical properties of eggs, which are critical for controlling nuclear size, are unique. In egg cytoplasm, abundant cytoplasmic molecules constitute numerous mesoscale condensates and increase the density of the molecules (Biswas *et al*., 2023; Keber *et al*., 2024). In particular, abundant RNAs in the egg cytoplasm can generate a hub for molecular condensates, such as nucleoli, through liquid-liquid phase separation and control the diffusivity of molecules depending on their charge state (Lafontaine *et al*., 2021; King *et al*., 2024). Therefore, these egg-specific molecular environments, with abundant RNAs, may contribute to the drastic changes in nuclear size, growth, and chromatin structure during early embryogenesis. Although RNAs in the egg cytoplasm are involved in controlling cellular processes, including nuclear and spindle assembly, by interacting with specific molecules (Grenfell *et al*., 2016; Aze *et al*., 2017), it remains unclear whether the RNA-based molecular environment of the egg cytoplasm contributes to the drastic conversion of nuclear structure after fertilization. To gain further insights, we evaluated the effects of abundant RNAs in *X. laevis* eggs in a cell-free reconstruction system, combined with the manipulation of RNA profiles.

## Results

### Degradation of RNAs reduces the growth speed of interphase nucleus size in a *Xenopus* egg extract

Total RNAs purified from unfertilized *X. laevis* eggs contained abundant ribosomal RNAs (rRNAs) (Fig. 1A), with a concentration range of approximately 8 mg/ml.

**Figure 1.**
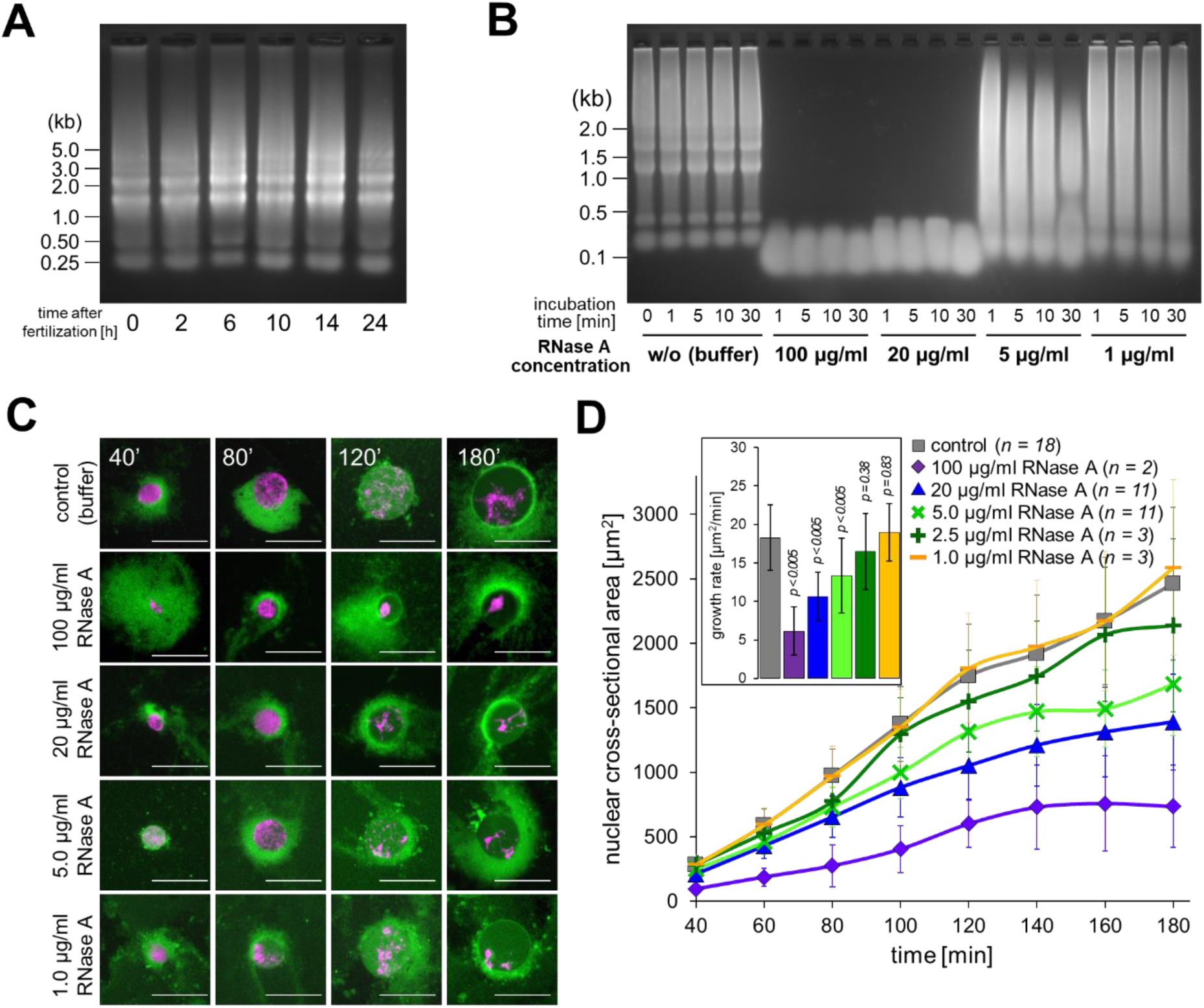
RNA degradation in the *X. laevis* egg extract. (**A**) The purified total RNAs from *X. laevis* unfertilized eggs and developed embryos at the indicated incubation period at 22°C after in vitro fertilization were analyzed by agarose gel electrophoresis. (**B**) The purified total RNAs from *X. laevis* egg extract supplemented with RNase A of the indicated concentration and incubated for the indicated duration were analyzed by agarose gel electrophoresis. The nucleotide sizes were shown based on the known DNA ladder on the left. (**C**) Representative images of the nuclei reconstructed in the *X. laevis* egg extract in the presence of RNase A of the indicated concentration and stained with Hoechst 33342 (for visualizing DNA; magenta) and DiOC_6_(3) (for visualizing membrane; green). Scale bars, 50 µm. (**D**) Dynamics of the mean cross-sectional areas of the nuclei reconstructed from *X. laevis* egg extract with each concentration of RNase A. Averages of mean cross-sectional areas (± SD) from each extract preparation at each incubation period are connected using a line in each condition with different RNase A concentration. The inset shows the average of calculated growth rates of nuclear cross-sectional area (± SD) from each extract preparation in each RNase A concentration. p values from Wilcoxon signed-rank test compared to the control condition were shown. Numbers (*n*) of experiments using each extract preparation for calculating the averaged growth rate were shown. The graph displaying the normalized values were shown in Fig. S1D.

During embryogenesis after fertilization, the profile of the purified total RNAs did not change substantially (Fig. 1A), which is consistent with experiments showing less transcriptional activity for major rRNA genes until the blastula stage (Shiokawa *et al*., 1979). To examine whether the abundant RNAs in the oocytes and developing embryos were involved in the rapid process of pronuclear assembly and size growth observed in fertilized eggs, we supplemented exonuclease RNase A into the cell-free reconstruction system from *X. laevis* egg cytoplasmic extract. In this cell-free system, after incubation with sperm chromatin for nuclear DNA material in the interphase extract, the nuclei were reconstructed within 40 min of incubation and grew continuously in size as incubation progressed, mimicking the in vivo male pronuclear assembly from sperm in cell-free conditions. After the addition of RNase A to the interphase egg extract, the purified total RNAs from the egg extracts were degraded into small RNAs (∼100 bp in length; Fig. 1B). The rate of RNA degradation was dependent on RNase A activity (Fig. 1B). These small residual RNAs were assumed to be protected from RNase degradation through interaction with other proteins, including ribosomal proteins (Ingolia *et al*., 2009) since the dissociation of ribosomal complexes is assumed to occur hours after incubation of the extract with RNase A (Choi *et al*., 2024). When sperm chromatin was incubated in the interphase extract in the presence of RNase A at different concentrations, the interphase nuclei were reconstructed within 40 min of incubation, similar to the control conditions (Fig. 1C). However, after the nuclear assembly phase, the growth rate of the observed nuclear cross-sectional area decreased in an RNase A concentration-dependent manner (Fig. 1D). We noted that the observed nuclear cross-sectional area varied among individual preparations of egg extracts, even under control conditions. Therefore, we set the criteria to allow us to use the measured data for analyses in this study (see Materials and Methods). We also confirmed the difference in the growth rate of the nucleus using normalized values of the nuclear cross-sectional area for each extract preparation (Fig. S1A). In addition to the reduction in nuclear growth speed, especially under conditions of higher RNase A concentration (such as > 20 µg/ml of RNase A), the chromatin within the reconstructed nuclei was more compacted after 80 min of incubation compared to the spread distribution of condensed chromatin within the nucleus under control conditions (Fig. 1C). The reduction in nuclear growth speed and compaction of chromatin was no longer observed when both RNase A and the RNase inhibitor were added to the egg extract (Fig. S1B–D), suggesting that the degradation of RNAs by RNase A, rather than the artificial effects of RNase A itself, causes changes in nuclear size and chromatin. The nuclei were not reconstructed from the sperm chromatin in the extract pre-incubated with RNase A before the addition of sperm chromatin, as previously shown (Fig. S1E; (Aze *et al*., 2017)). It is suggested that the intact RNAs present before degradation by RNase A are involved in the reconstruction of nuclei in the presence of RNase A (Fig. 1C).

These data suggest that the abundant RNAs in the egg extract facilitate the rapid growth of the interphase nucleus and disperse the distribution of chromatin within the nucleus.

### Optimal concentration of RNAs maximizes growth rate of nuclear size

To determine the most effective concentration and size of RNAs for interphase nuclear size growth, we manipulated the concentration and profile of RNAs by adding extra RNAs to the egg extract. First, the total RNAs purified from the egg extract (hereafter referred to as “native RNAs”) were added to the egg extract with sperm chromatin (Fig. 2A). When the RNA concentration was increased by 0.1 mg/ml by adding native RNAs to the egg extract, the resulting growth rate of nuclear size increased slightly compared to the control condition (Fig. 2A, B). In contrast, in the condition adding extra native RNAs greater than 0.5 mg/ml, the resulting growth rate of the nucleus was reduced (Fig. 2A, B). Furthermore, the chromatin within the reconstructed nuclei showed a more dispersed distribution after 80 min of incubation than under control conditions (Fig. 2A). These data suggest that the RNA concentration in conventional egg extracts is not the most effective for accelerating interphase nuclear size growth. While a smaller amount of additional RNAs can facilitate growth, excessive amounts of RNAs inhibit rapid nuclear size growth.

**Figure 2.**
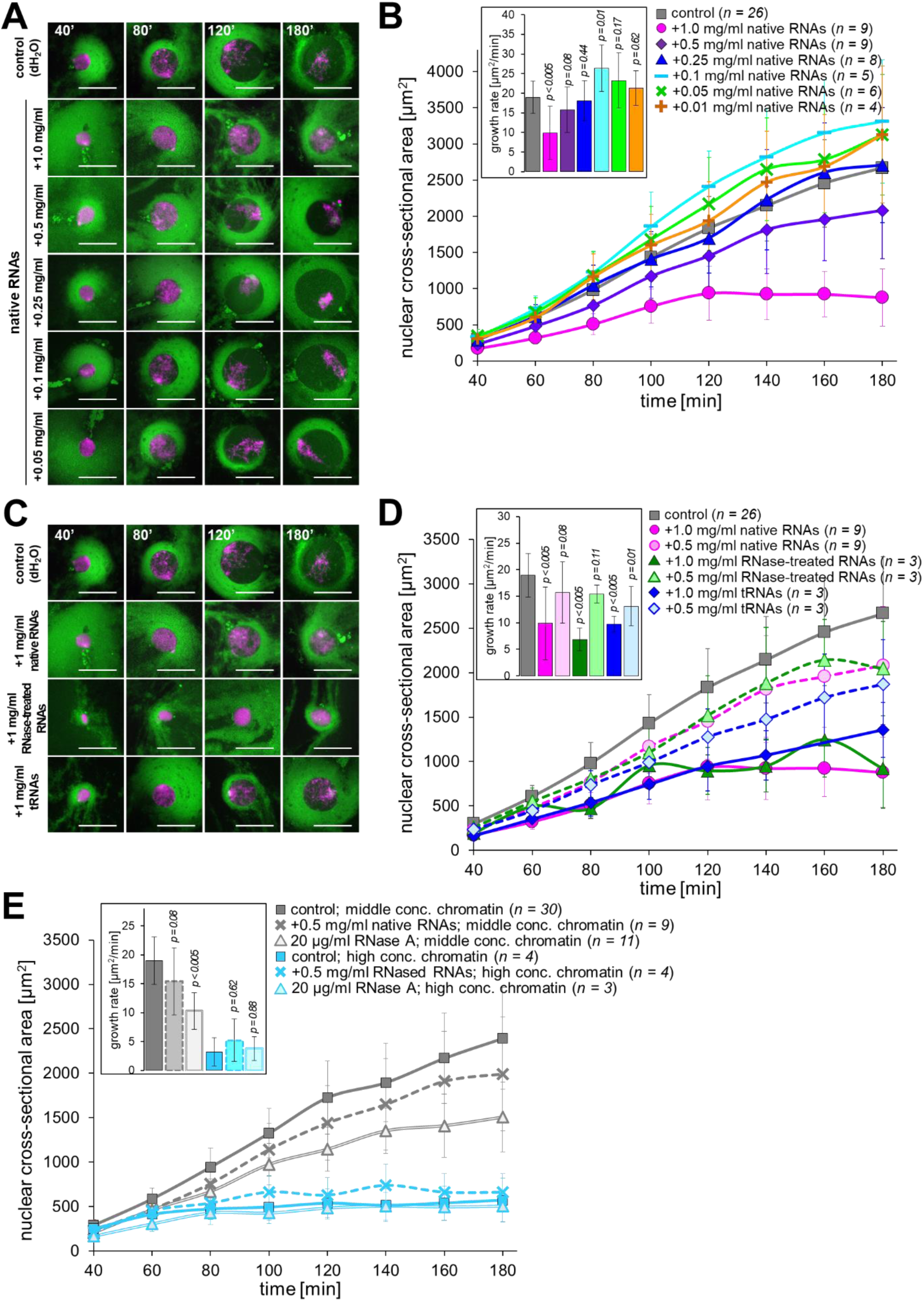
Supplementation of RNAs to the *X. laevis* egg extract. (**A**) Representative images of the nuclei reconstructed from the *X. laevis* egg extract by supplementation of the native RNAs at different concentrations and stained with Hoechst 33342 (for visualizing DNA; magenta) and DiOC_6_(3) (for visualizing membrane; green). Scale bars, 50 µm. (**B**) Dynamics of the mean cross-sectional areas of the nuclei reconstructed from *X. laevis* egg extract in the presence of native RNAs at the indicated concentration. Averages of mean cross-sectional areas (± SD) from each extract preparation are connected using a line in each dataset. Numbers (*n*) of experiments using each extract preparation for calculating averaged growth rate were shown. The graph displaying the normalized values were shown in Fig. S2A. (**C**) Representative images of the nuclei reconstructed from the *X. laevis* egg extract in the presence of each type of RNAs (native RNAs, RNase-treated RNAs, and tRNAs) at additional 1.0 mg/ml and stained with Hoechst 33342 and DiOC_6_(3). Scale bars, 50 µm. (**D**) Dynamics of the mean cross-sectional areas of the nuclei reconstructed from *X. laevis* egg extract in the presence of each type of RNAs at either concentration 0.5 or 1 mg/ml. The inset shows the average of calculated speeds of nuclear cross-sectional area (± SD) in each condition with p values from the Wilcoxon signed-rank test compared to the control condition. The graph displaying the normalized values were shown in Fig. S2B. (**E**) Dynamics of the mean cross-sectional areas of the nuclei reconstructed from different concentrations of sperm chromatin (normal conc.: ∼300 sperm/µl, grey; high conc.: ∼5,000 sperm/µl, light blue) incubated in the *X. laevis* egg extract in the presence of native RNAs at additional 1.0 mg/ml or RNase A at 20 µg/ml. The graphs displaying the normalized values by each extract preparation were shown in Fig. S2C and S2D.

Considering that the effective range of added concentration (0.1–1 mg/ml) on the nuclear size growth is comparable to the 1.2–12% increase in the total RNA concentration in the egg extract, the functional amount of RNAs for controlling the nuclear size growth is assumed to be much less than the superficial amount of total RNAs (∼8 mg/ml) from the egg extract. The majority of endogenous RNAs that interact with ribosomal proteins and are protected from RNA degradation by RNase A are not expected to be involved in controlling nuclear growth. To examine the effects of the RNA profiles, we added different types of RNAs to the egg extract.

When the total RNAs purified from the egg extract incubated with RNase A (hereafter as “RNase-treated RNAs”) or yeast tRNAs (“tRNAs”) were added at relatively higher concentrations of 0.5 or 1 mg/ml to the egg extract, the resulting growth rate of the nucleus was reduced compared to the control conditions and showed the dependence on the added RNA concentration (Fig. 2C, D). Furthermore, the chromatin within the reconstructed nuclei showed a more dispersed distribution, especially in the presence of higher concentrations of either type of RNA (Fig. 2C).

Although the effects of RNAs on reducing the nuclear growth rate showed slight differences among the types of RNAs added, the amount of RNAs in the egg extract could regulate the nuclear size and growth rate, regardless of the type of RNAs.

These results suggest that the relative amount of RNAs per nucleus, rather than the absolute amount or concentration in the egg extract, may contribute to the control of nuclear growth. To examine this possibility, we manipulated the concentration of sperm chromatin in the egg extracts (Fig. 2E). Under the condition of a higher concentration of sperm chromatin (∼5,000 sperm/µl), nuclei grew slower in size even without excessive RNAs or RNase A compared to the conventional concentration (∼300 sperm/µl) condition, as observed previously (Hara and Merten, 2015). When incubated with excessive native RNAs (+ 0.5 mg/ml) or RNase A (20 µg/ml) in the higher sperm concentration condition, the changes in the speed of nuclear size growth appeared to be less compared to those under the conventional sperm concentration condition (Fig. 2E). In particular, when excessive RNAs were added to the extract containing a higher concentration of sperm chromatin, the resulting speed of nuclear size growth increased, which is the opposite of the observed reduction in nuclear size growth speed under conventional sperm concentration conditions (Fig. 2E). When calculating the relative amount of RNAs per sperm, the change in the relative amount of RNAs is + ∼0.1 ng/sperm; from 1.6 ng/sperm (8 µg of RNA/5,000 sperm) in the control condition to 1.7 ng/sperm (8.5 µg/5,000 sperm) after addition of RNAs plus 0.5 mg/ml to the egg extract. This change is lower than that in the conventional sperm concentration condition (+ ∼1.7 ng/sperm; from 26.6 ng/sperm [8 µg of RNA/300 sperm] to 28.3 ng/sperm [8.5 µg/300 sperm]) and comparable to that observed in the rapid growth of nuclear size by adding 0.1 mg/ml of RNAs to the egg extract containing the conventional sperm concentration condition (+ ∼0.3 ng/sperm; from 26.6 ng/sperm [8 µg of RNA/300 sperm] to 27 ng/sperm [8.1 µg/300 sperm]; Fig. 2B). These results suggest that the optimal amount of RNAs depends on the number of nuclei in egg extracts.

### Cytoplasmic RNAs facilitate the primary step of nuclear assembly

To explore how RNAs regulate nuclear growth, we first analyzed the cellular localization of RNAs. When the egg extract containing the reconstructed nuclei was stained with dyes specific for the detection of RNAs, the stained signal was rarely detected within the nuclei, even when either RNA-specific dye was used (Fig. 3A). Consistent with the reduced localization of RNAs within the nuclei, the nucleolar structures, which generally form puncta containing rRNAs, were rarely detected as puncta within the reconstructed nuclei by immunostaining for the nucleolar proteins fibrillarin and nucleophosmin (Fig. 3A). The detected weak signals suggested that the premature state of nucleoli could be formed within the nuclei reconstructed from *X. laevis* egg extract, as shown previously in early mouse embryos (Kresoja-Rakic and Santoro, 2019). In contrast, in the nuclei isolated from developed embryos at the gastrula stage, RNAs and nucleolar proteins were detected as puncta within the nuclei (Fig. 3B). Thus, RNAs and nucleolar structures were less localized within the nuclei reconstructed from *X. laevis* egg extracts, and putatively in early stage embryos.

**Figure 3.**
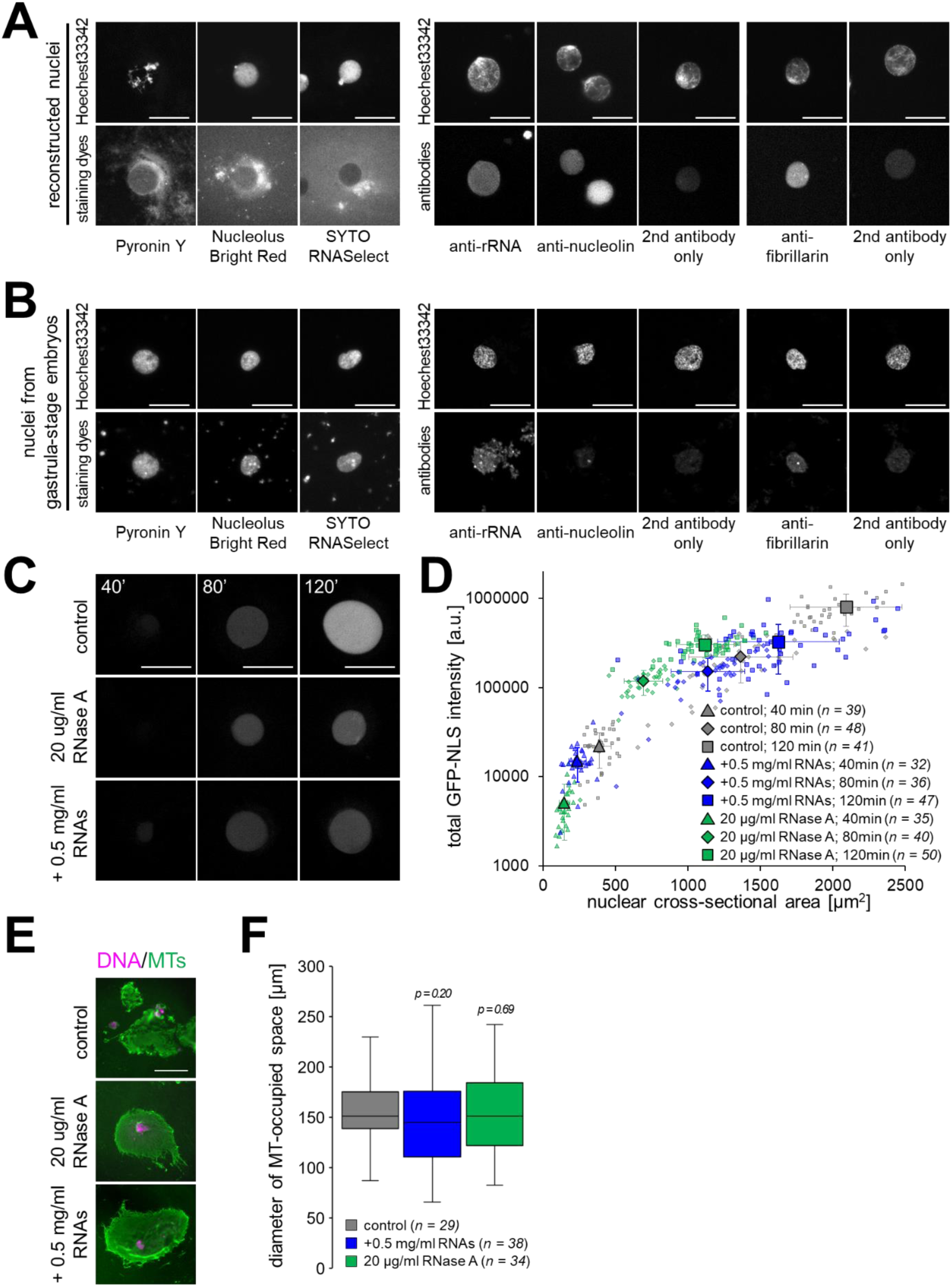
Intracellular localization of RNAs and contribution of cytoplasmic mechanisms for nuclear size control. (**A**) Representative images of the nuclei reconstructed from the *X. laevis* egg extract and stained with Hoechst 33342 and RNA staining dyes (Pyronin Y, Nucleolus Bright Red, SYTO RNASelect) or immunostained using anti-nucleolin or fibrillarin antibodies in the sedimented samples on a glass slide. The images from immunostaining using only secondary antibodies were also shown. Scale bars, 50 µm. (**B**) Representative images of the nuclei isolated from developed embryos at gastrula stage stained by RNA staining dye or immunostained for nucleolar proteins. Scale bars, 20 µm. (**C**) Representative images of the nuclei reconstructed from the *X. laevis* egg extract containing GFP-NLS recombinant proteins and additional native RNAs at 0.5 mg/ml or 20 µg/ml RNase A at indicated time of incubation. Scale bars, 50 µm. (**D**) Scatter plots of the GFP-NLS intensity per observed nuclear cross-sectional area with the measured nuclear cross-sectional area among samples reconstructed with different conditions at each incubation time. Large and small symbols represent the mean values (±SD) among the samples and obtained values of the individual nucleus, respectively. (**E**) Representative images of the nuclei reconstructed from the *X. laevis* egg extract and stained using α-tubulin antibody (green; immunostaining for visualizing microtubules (MTs)) and Hoechst 33342 (magenta; staining for DNA) in the sedimented samples on a glass slide. Scale bar: 100 μm. (**F**) Measured mean diameter (± SD) of MT-occupied space in each sample upon supplementation with additional native RNAs or RNase A. p values from the Wilcoxon test compared to the control condition were shown.

Control of nuclear size growth involves the supply of nuclear membrane components, including nuclear proteins and membranes, via the nuclear import machinery and supply from the proximal endoplasmic reticulum, respectively. To evaluate the involvement of RNAs in these cytoplasmic processes, we first analyzed nuclear import activity by visualizing the GFP-fused nuclear localization signal peptide (GFP-NLS protein), which can be imported into the nucleus. Quantification of the signal intensity of the imported GFP-NLS proteins in the nuclei showed that the intensity was lower in the presence of RNase A or an excess amount of native RNAs (Fig. 3C, S3A). This slow import of GFP-NLS proteins can explain the delay in DNA replication rate in conditions treated with RNase A or excess amounts of RNAs (Fig. 3SB; (Aze *et al*., 2017)) due to the putative slow import of machinery proteins for DNA replication into the nucleus. However, the calculated total intensities of GFP-NLS in nuclei incubated with RNase A and excess RNAs were comparable to those of the same nuclear size in the early period of incubation under control conditions (Fig. 3D). This suggests that nuclear import activity, which spontaneously correlates with nuclear size, is not drastically altered by manipulating the amount of RNAs in the extract. We noted that the slow increase in DNA quantity due to slow DNA duplication may explain the observed slow growth in nuclear size, based on the known DNA quantity-dependent control of nuclear size growth (Heijo *et al*., 2020).

When RNase A was added to the egg extract containing the DNA replication inhibitor aphidicolin, slow growth of the nucleus was still observed compared to that in the absence of RNase A (Fig. S3C), suggesting that the RNA-based mechanisms controlling nuclear growth are mainly independent of DNA quantity-dependent mechanisms. Next, we measured the size of the microtubule space around the reconstructed nucleus, which is known to correlate with the growth rate of the nucleus (Hara and Merten, 2015; Mukherjee *et al*., 2020) in the presence of RNase A or extra RNAs. The measured sizes of the immunostained microtubule-occupied space were not significantly different in the presence of RNase A or excess RNAs (Fig. 3E, F). These results suggest that the amount of RNAs in egg extract does not substantially influence cytoplasmic-based mechanisms for nuclear growth.

Considering the observed lower contribution of cytoplasmic RNAs to the known cytoplasmic-based mechanisms, it is possible that RNAs contribute to the primary step of nuclear assembly before the nuclear membrane covers the chromatin, rather than to the later step of nuclear growth after nuclear envelope formation. To distinguish the effects of RNAs on the later nuclear growth step from those on the early nuclear assembly step, an excess amount of RNAs or RNase A was added to the egg extract containing nuclei after 60 min of incubation with sperm chromatin to complete DNA replication (Fig. 4A, B). In the presence of RNAs or RNase A, the nuclei continued to grow in size for an additional 60 min of incubation, as observed under the control condition, and displayed a similar chromatin distribution within the nucleus (Fig. 4A, B). However, after an additional 60 min of incubation, the growth rate of the nucleus was slightly reduced compared with the control condition (Fig. 4B). These data suggest that the effectiveness of RNAs in controlling nuclear growth is attenuated after nuclear assembly in interphase.

**Figure 4.**
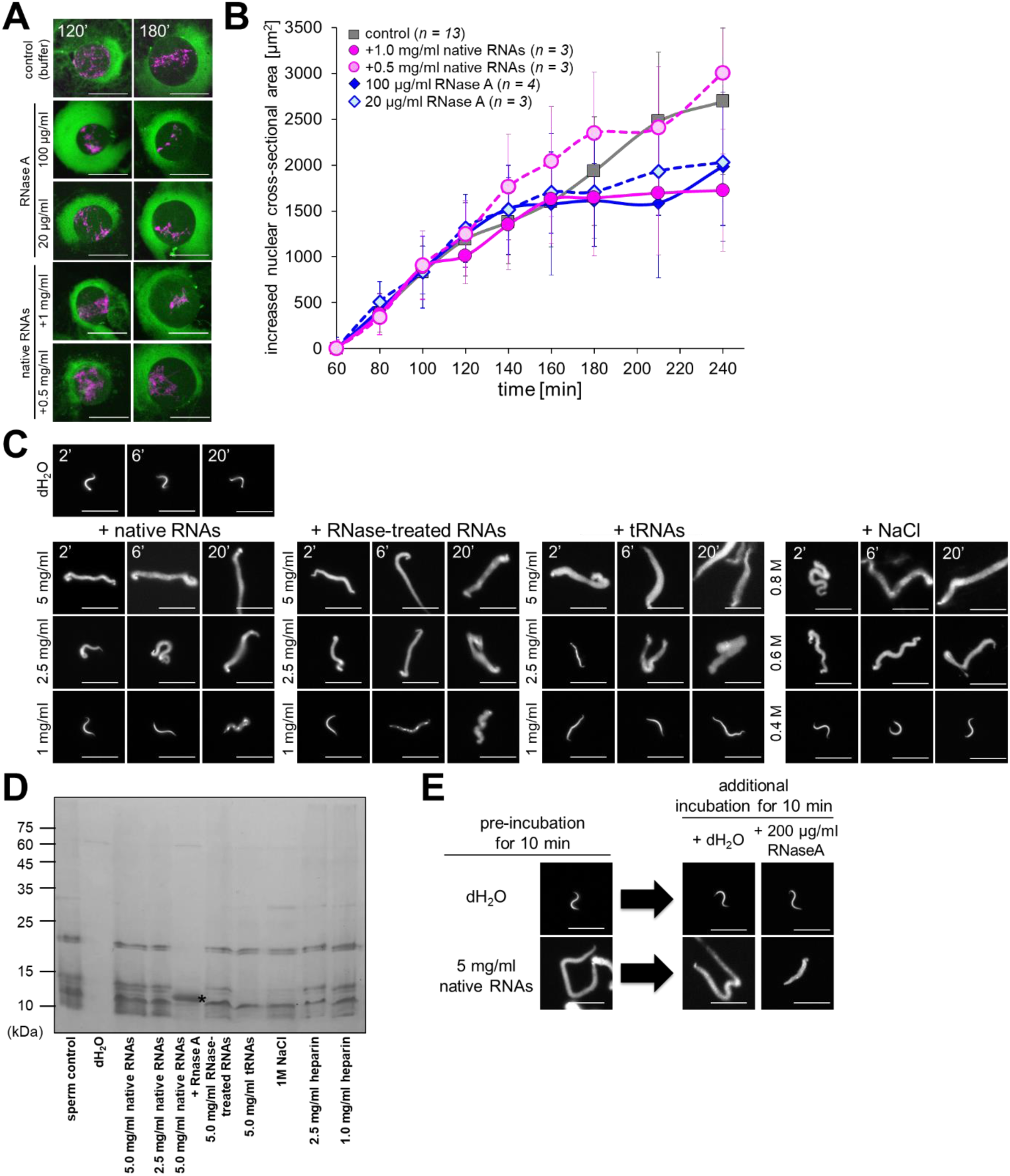
RNAs facilitate chromatin decompaction before covering nuclear membrane. (**A**) Representative images of the nuclei reconstructed from the *X. laevis* egg extract by supplementation of RNase A or native RNAs at the indicated concentration after 60 min of incubation and stained with Hoechst 33342 (for visualizing DNA; magenta) and DiOC_6_(3) (for visualizing membrane; green). The total time of incubation with sperm chromatin in the egg extract was shown. Scale bars, 50 µm. (**B**) Dynamics of the increased cross-sectional areas of the nuclei reconstructed from *X. laevis* egg extract by supplementation of RNase A or native RNAs at the indicated concentration after 60 min of incubation. The increased nuclear cross-sectional area was calculated as the mean cross-sectional areas at 60 min of incubation before the supplementation of RNAs or RNase A were subtracted from the mean areas at each incubation time. Averages of the mean increased areas (± SD) from each extract preparation are connected using a line in each dataset. Numbers (*n*) of experiments using each extract preparation were shown. The graph displaying the normalized nuclear cross-sectional areas by each extract preparation were shown in Fig. S4A. (**C**) Representative images of the sperm chromatin incubated with solution containing each type of RNAs or NaCl at indicated concentration for the indicated time. The chromatin was stained with Hoechst 33342. Scale bars, 20 µm. (**D**) Protein profile of the supernatant after incubation of sperm chromatin with solution containing RNAs (native RNAs, RNase-treated RNA, tRNA) or ionic compounds (heparin, NaCl) at the indicated concentrations for 10 min analyzed by CBB-staining of SDS-PAGE gel. The most left lane represents the protein profile from the whole sperm chromatin as control. * shows the band relative to the RNase A which was added after preincubation of sperm chromatin. (**E**) Representative images of sperm chromatin incubated by adding 200 µg/ml of RNase A for 10 min after preincubation with 5 mg/ml of native RNAs.

To further analyze the effects of RNAs on the primary step of nuclear assembly, sperm chromatin was incubated directly in a solution containing RNAs without egg extract. After a few minutes of incubation with RNAs, the chromatin swelled and converted to the elongated shape of the original sperm, regardless of the type of RNAs used (native RNAs, RNase-treated RNAs, and tRNAs) (Fig. 4C). The intensity of chromatin staining with the DNA-specific dye decreased in RNA-treated chromatin, indicating that the chromatin was dispersed and decompacted. The degree of decompaction progressed with increasing incubation period and showed a concentration dependence of RNAs (Fig. 4C). In addition to morphological changes in chromatin, some proteins were detected in the supernatant of sperm chromatin incubated with any of the RNA types (Fig. 4D). The protein pattern detected after incubation with RNAs was similar to that observed after incubation with a high concentration of NaCl (Fig. 4D). Since NaCl treatment induces the decompaction of sperm chromatin and promotes the dissociation of nuclear proteins, including histones and protamines, from sperm into the supernatant (Ohsumi and Katagiri, 1991), RNAs are predicted to dissociate protamines and histones from sperm chromatin, even in the absence of the chaperone protein nucleoplasmins, which facilitates the removal of protamines and the incorporation of histones into chromatin. Interestingly, the decompacted chromatin after incubation with native RNAs was recondensed upon RNA degradation by the addition of RNase A (Fig. 4E), and most of the dissociated proteins were no longer detected in the supernatant (Fig. 4D), although the size of the resulting chromatin appeared to be slightly increased compared to the original sperm chromatin. These data suggest that RNAs surrounding sperm chromatin are involved not only in the dissociation of nuclear proteins from sperm chromatin, but also in the decompaction of chromatin during the primary step of interphase nuclear assembly.

### Negative charges of RNAs facilitate decompaction of sperm chromatin

Considering that the nucleotide sequences themselves are less important for controlling the nuclear growth speed (Fig. 2) and sperm chromatin decompaction (Fig. 4), we focused on the fundamental property of RNAs, the negative charge of RNAs in solution. To study the effects of negative charges on chromatin decompaction, we incubated sperm chromatin with charged compounds, such as negatively charged heparin or positively charged protamine from salmon sperm. In the presence of heparin, sperm chromatin decompacted in a concentration-dependent manner (Fig. 5A), and proteins dissociated from sperm chromatin (Fig. 4D). In contrast, sperm chromatin was not morphologically altered in the presence of protamines (Fig. 5A). Furthermore, chromatin that was decompacted by incubation with either native RNAs or heparin recondensed and showed an aggregated state upon the addition of positively charged protamines (Fig. 5B). These results suggest that the observed reversible changes in the compaction state of sperm chromatin by RNAs were reproduced using charged molecules. Next, to assess whether the observed effects of negative charges were limited to sperm chromatin, we incubated nuclei isolated from developed embryos with solutions containing charged compounds or RNAs (Fig. 5C). Although incubation with heparin or NaCl caused the chromatin to spread out from the nucleus, forming a halo-like structure, incubation with native RNAs did not induce the decompaction of chromatin within the nucleus, even at the highest concentration that could induce decompaction when sperm chromatin was used (Fig. 5C). The lower sensitivity of RNAs to chromatin decompaction in embryonic nuclei compared to sperm chromatin may be due to differences in the accessibility of native RNAs from the cytoplasm to chromatin. RNA molecules, which are long polymers, are not diffusely imported through the nuclear pores after the formation of the nuclear envelope, as evidenced by the reduced localization of RNAs within the reconstructed nuclei (Fig. 3A), whereas RNAs are accessible to bare chromatin before the nuclear membrane covers the chromatin.

**Figure 5.**
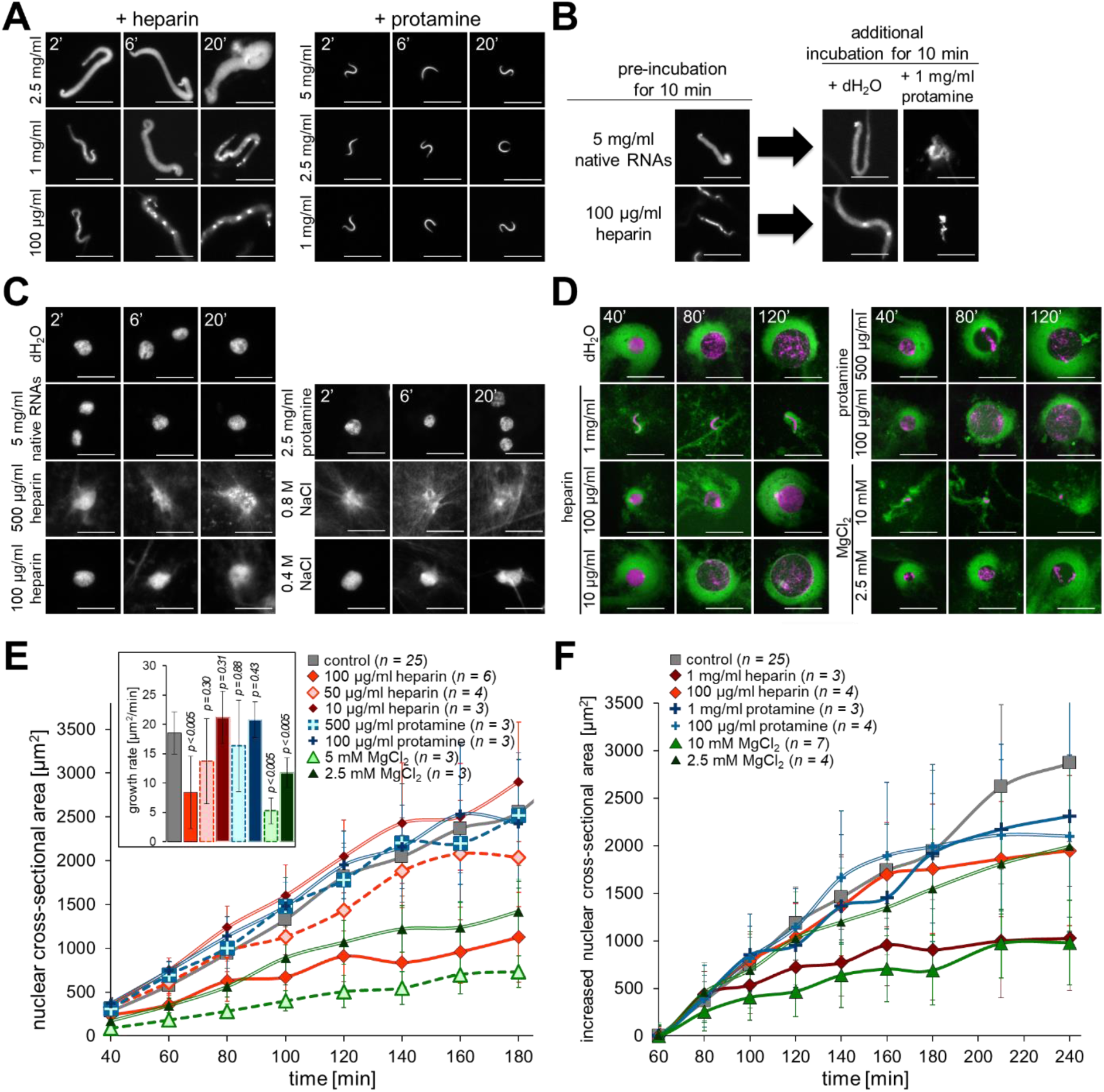
Negatively charged molecules facilitate decompaction of sperm chromatin and nuclear growth. (**A**) Representative images of the sperm chromatin incubated with solution containing negatively charged heparin, positively charged protamine, or NaCl at the indicated concentration for the indicated time. The chromatin was stained with Hoechst 33342. Scale bars, 20 µm. (**B**) Representative images of the sperm chromatin incubated by adding 1 mg/ml protamine solution for 10 min after preincubation with 5 mg/ml native RNAs or 100 µg/ml heparin solution. (**C**) Representative images of the nuclei isolated from developed embryos at the gastrula stage after incubation with native RNAs, ionic solution containing heparin, protamine, or NaCl at the indicated concentration. The chromatin was stained with Hoechst 33342. (**D**) Representative images of the nuclei reconstructed from *X. laevis* egg extract by adding solution containing heparin, protamine, or MgCl_2_ at different concentrations. The extract containing the nuclei was stained with Hoechst 33342 (magenta) and DiOC_6_(3) (for visualizing membrane; green). Scale bars, 50 µm. (**E**) Dynamics of the mean cross-sectional areas of the nuclei reconstructed from *X. laevis* egg extract in the presence of each ionic solution. Averages of mean cross-sectional areas (± SD) from each extract preparation are connected using a line in each dataset. The inset shows the average of calculated growth rates of nuclear cross-sectional area (± SD) in each condition with p values from the Wilcoxon test compared to the control condition. Numbers (*n*) of experiments using each extract preparation for calculating the averaged growth rate were shown. The graph displaying the normalized values by each extract preparation were shown in Fig. S5A. (**F**) Dynamics of the increased cross-sectional areas of the reconstructed nuclei from *X. laevis* egg extract by supplementation of each ionic solution at the indicated concentration after 60 min of incubation with sperm chromatin. The increased nuclear area was calculated as the mean cross-sectional areas at 60 min of incubation before the supplementation of ionic solution were subtracted from the mean areas at each incubation time points. The graph displaying the normalized values by each extract preparation were shown in Fig. S5B.

Next, to examine the contribution of negatively charged RNAs to interphase nuclear growth, charged compounds were added to the egg extract. First, negatively charged heparin or positively charged protamines were added to the egg extract simultaneously with the sperm chromatin (Fig. 5D, E). In the presence of heparin at a relatively low concentration (10 µg/ml), the resulting growth speed of nuclear size was slightly increased compared to the control condition (Fig. 5A, B). In contrast, in the presence of heparin at a higher concentration (> 50 µg/ml), the chromatin within the nucleus showed a dispersed distribution throughout the nucleus, and the resulting growth speed of nuclear size was reduced in a concentration-dependent manner. At the highest concentration (1 mg/ml), the chromatin did not change from the sperm shape during the incubation period (Fig. 5D). These concentration-dependent features of nuclear growth rate and chromatin condensation status were similar to those of RNA supplementation (Fig. 2). Furthermore, the reduction in the growth rate of nucleus was recapitulated when MgCl_2_ was added, as shown previously (Heijo *et al*., 2020). However, the resulting chromatin was more condensed in the aggregated state, which is unlikely in condensed chromatin with an evenly distributed pattern in the nuclei supplemented with heparin or RNAs (Fig. 5D, E). Mg^2+^ cations neutralize the negative charge of chromatin, resulting in excessive chromatin compaction and reduced growth rateof the nucleus (Shimamoto *et al*., 2017; Maeshima *et al*., 2018; Heijo *et al*., 2020). These discrepancies in chromatin distribution suggest that the regulatory pathways to attain the reduction in nuclear size growth are different when using positively charged Mg^2+^ and negatively charged RNAs or heparin. When these ionic compounds were added to the extract after nuclear assembly with complete DNA replication, the time point showing slow nuclear growth in conditions supplemented with heparin or MgCl_2_ (Fig. 5E) was much earlier than that of supplementation with RNAs (Fig. 4). Native RNAs are predicted to be polymers of nucleotides that are too long to diffuse through nuclear pores, whereas much smaller Mg^2+^ ions or putative small polymer heparins can diffuse, suggesting that the difference in the accessibility of the molecules to the chromatin inside the nucleus could result in different condensation states and nuclear growth speeds.

### Negative charges of RNAs facilitate the deposition of nucleosome proteins in chromatin

Increased amounts of either anionic or cationic molecules induced a reduction in the growth rate of the nucleus. To distinguish the regulatory pathways regarding the charge status, nucleosome protein histones were immunostained in the reconstructed nuclei under conditions of supplementation with a solution containing negatively or positively charged compounds in the egg extract. In the control condition, histone proteins, such as histone H3 or histone H2B, which are incorporated into the sperm chromatin from the cytoplasm, were detected in the chromatin of the reconstructed nuclei (Fig. 6A). In the presence of RNase A in the egg extract, the amount of histones detected on the chromatin in the reconstructed nuclei was drastically reduced (Fig. 6A–C). In contrast, when heparin or native RNAs was added at a higher concentration, at which the nuclear growth rate was reduced, more histones were incorporated into the chromatin than in the control condition (Fig. 6A–C). Considering that heparin and RNAs facilitate the removal of nuclear proteins from sperm chromatin (Figs. 4 and 5), these experimental results suggest that the remodeled chromatin with fewer nuclear proteins, in turn, facilitates the incorporation of free histones into nuclear chromatin, presumably via the histone chaperone nucleoplasmin present in the egg extract. Similarly, when positively charged protamines were added to egg extract, the amount of incorporated histones was significantly reduced (Fig. 6B, C). The added protamines may prevent the removal of existing protamines from sperm chromatin, resulting in a fewer amount of newly incorporated histones. However, when MgCl_2_, another anion Mg^2+^ that induces chromatin compaction and reduces nuclear growth rate, was added to the extract, the changes in the incorporated histones were less pronounced compared to the conditions with the addition of heparin, RNAs, or protamine (Fig. 6B, C). This suggests that Mg^2+^ anions preferentially affect the excessive condensation of chromatin by neutralizing the electrical repulsion of negatively charged chromatin rather than the deposition of nucleosome proteins. To further evaluate whether these observed differences in histone composition in the chromatin of the reconstructed nuclei could contribute to the observed differences in nuclear growth rate, we compared the quantified amount of histones in the chromatin region with the observed nuclear cross-sectional area after a given incubation time. The nuclear cross-sectional area is known to correlate with the growth rate of nuclear size in cell-free reconstruction systems (Hara and Merten, 2015). From the scatter plot of these two parameters from all measured data under different conditions, the histone amount correlated very weakly with the nuclear size, not only among data from individual nuclei with intrinsic size variation in the single experimental condition (Fig. S6A) but also among the averaged data from all experimental conditions (Fig. 6D; R^2^ = 0.32). In particular, although samples incubated with RNase A or excess RNAs showed a smaller reconstructed nucleus than the control condition, the amount of incorporated histones in the presence of excess RNAs was significantly higher than that in the presence of RNase A. Instead, when we calculated the entropy of the chromatin within the nucleus, which is a parameter related to the distribution of chromatin (Heijo *et al*., 2020), the entropy value correlated better with the nuclear size among data from individual nuclei and averaged values from each condition (Fig. 6E; R^2^ = 0.78; Fig. S6B). We noted that the entropy value also correlated with the amount of histones incorporated into the chromatin (Fig. 6F; R^2^ = 0.73), suggesting that the nucleosome composition of chromatin within the nucleus, which is regulated by ionic conditions around the chromatin, indirectly contributes to the regulation of nuclear growth speed by altering chromatin distribution and structure. Finally, when data on incorporated histones, entropy, and nuclear size of the isolated nuclei from developed embryos were obtained, the amount of histones in the nuclei was significantly lower than that in the reconstructed nuclei from unfertilized eggs (Fig. 6D). In contrast, the data from the developed nuclei plotted around the regression line between entropy and nuclear size from the data of the reconstructed nuclei (Fig. 6E). The correlation between chromosome distribution and nuclear size can explain the data obtained from our cell-free reconstructed nuclei and the isolated in vivo nuclei of the developed embryos.

**Figure 6.**
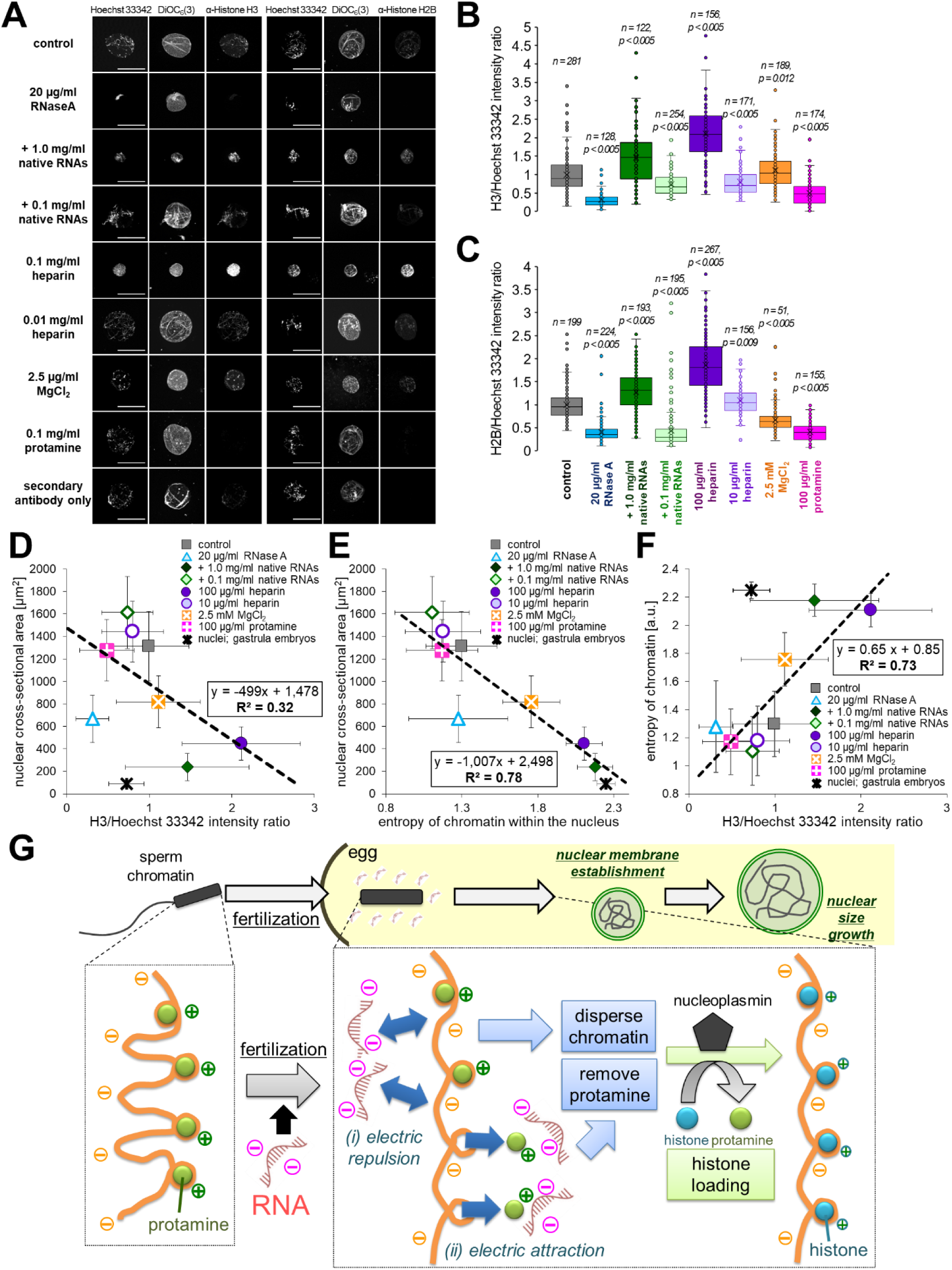
RNAs facilitate deposition of nucleosome proteins. (**A**) Representative images of the reconstructed nuclei immunostained using anti-histone H3 or histone H2B antibodies. The nuclei were reconstructed in the *X. laevis* egg extract in the presence of the indicated supplements for 120 min of incubation with sperm chromatin. The immunostained images using only secondary antibodies were also shown. Scale bars, 50 µm. (**B, C**) Box plots of the intensities quantified from the chromatin within the nucleus immunostained using anti-histone H3 (**B**) or anti-histone H2B (**C**). The calculated intensity ratio of histone/Hoechst 33342, dividing the measured total intensity using each antibody throughout the nuclear cross-sectional area by the total intensity stained with Hoechst 33342, was normalized by the mean intensity ratio obtained from the control condition. Line within the box represents the median. The upper and bottom lines represent the 75th and 25th percentile. The whiskers represent the 95th or 5th percentile. The individual data was shown as the circle symbol. p values from the Student’s t-test compared to the control condition and numbers (*n*) of quantification in each condition were shown. (**D**) Scatter plots of the measured mean nuclear cross-sectional areas in each condition of the indicated supplements with the intensity ratio of histone H3/Hoechst 33342. (**E**) Scatter plots of the measured mean nuclear cross-sectional areas in each condition of the indicated supplements with the mean calculated value of the chromatin entropy. (**F**) Scatter plots of the calculated value of the chromatin entropy in each condition of the indicated supplementation with the relative intensity ratio of histone H3/Hoechst 33342. Plots showing the individual data of each nucleus regarding (**D, E**) were shown in Fig. S6A and S6B. Plots regarding (**D, F**) by using H2B data were shown in Fig. S6C and S6D. Dataset from the nuclei reconstructed in the egg extract (colored symbols) was fitted using a linear regression with showing the equation and the value of determination of the correlation (R^2^). Additionally, the data from the isolated nuclei from embryos at gastrula stages (black symbol) was plotted. (**G**) Putative mechanisms how RNAs contribute to the chromatin decompaction after fertilization. The highly condensed chromatin with wrapping to protamines in sperm initiates to interact with cytoplasmic RNAs in the egg cytoplasm after fertilization. (i) The negative charges of RNAs generate the repulsive force against the negatively charged chromatin, resulting in the conversion of chromatin structure to dispersed state. Simultaneously, (*ii*) negative charges of RNAs attract to positive charges of protamines and/or nuclear proteins, resulting in the detachment of the nuclear proteins from the sperm chromatin. These decompacted chromatin with less protamines facilitates the incorporation of new histones from the egg cytoplasm via nucleoplasmin. This conversion of the sperm chromatin helps the pronuclear assembly become more rapid and the growth speed of the interphase nuclear size increase.

## Discussion

### RNAs are involved in the rapid assembly of pronuclei after fertilization in *Xenopus* eggs

RNAs, primarily rRNAs, are abundant in *X. laevis* eggs. rRNAs serve as structural components of ribosomal complexes that facilitate protein translation in the cytoplasm. In addition to their general role in translation, recent studies have shown that RNAs contribute to the establishment of the intracellular molecular environment. rRNA interacts physically with nucleolar proteins through RNA binding and acidic motifs, leading to the formation of molecular condensates such as the nucleolus, via liquid-liquid phase separation (Lafontaine *et al*., 2021; King *et al*., 2024).

Furthermore, the negative charges of RNAs regulate the diffusivity of positively charged molecules in the cytoplasm of *X. laevis* eggs (Choi *et al*., 2024). Our cell-free reconstruction experiments provide new evidence that negatively charged RNAs in fertilized *X. laevis* eggs contribute to the rapid decompaction of sperm chromatin, which is then remodeled into round embryonic pronucleus. While the ionic environment contributes to the decompaction of sperm chromatin by incubating ionic molecules with sperm chromatin (Philpott *et al*., 1991), the molecular candidates that set the charge state in eggs have not been identified. Additionally, when the histone chaperone nucleoplasmins, known for their role in removing positively charged protamines and incorporating histones into chromatin (Ohsumi and Katagiri, 1991; Philpott and Leno, 1992), were inhibited in egg extract, chromatin decompaction was still observed (Biswas *et al*., 2023). In fertilized eggs from mice with a knockout of the nucleoplasmin gene NPM2, chromatin remains decondensed and forms a normal-shaped male pronucleus, although with a slightly altered chromatin distribution pattern (Burns *et al*., 2003). This evidence supports the involvement of other nucleoplasmin-independent molecules in sperm chromatin decompaction. RNA degradation by pre-incubating the extract with RNase A (Fig. S1E; (Aze *et al*., 2017)) or the addition of excessive amounts of heparin (Fig. 5D) completely inhibited the decompaction of sperm chromatin, supporting the primary involvement of cellular RNAs in this process. However, since most cytoplasmic proteins are negatively charged (Wühr *et al*., 2014), it is possible that other negatively charged proteins, in addition to RNAs, may partially contribute to the rapid pronuclear assembly.

Moreover, as RNAs degrade, polymeric RNAs and RNA-containing protein complexes dissociate into numerous monomeric nucleotides and individual components, which can increase cytoplasmic osmolality by generating more osmolytes. The rise in osmolality within the nucleus or the condensed region of chromatin facilitates the flux of small molecules, contributing to an increase in nuclear size and decompaction of chromatin (Mitchison, 2019; Wu *et al*., 2022; Hara, 2023). Contrary to the expected osmotic response, the addition of RNase A after nuclear assembly, which is thought to increase nucleotides and dissociated proteins in the cytoplasm, did not induce an acute response in nuclear size or chromatin distribution (Fig. 4). This result complicates the explanation of the changes in nuclear size and chromatin distribution observed in this study solely through osmolarity-based mechanisms. During embryogenesis, although the detected concentration and profile of RNAs from embryos did not change drastically (Fig. 1A), the relative amount of RNAs per cell, nucleus, or genomic DNA was greatly reduced because the cell volume is almost halved at each embryonic cleavage. Furthermore, the amount of histone chaperone nucleoplasmin in cells decreases during embryogenesis (Chen *et al*., 2019). Therefore, the contribution of both RNAs and nucleoplasmin to chromatin remodeling may decrease as the embryo develops. In line with this, we observed an increase in intranuclear RNAs and nucleolar proteins during embryogenesis (Fig. 3), consistent with previous reports (Arai *et al*., 2017; Ke *et al*., 2017; Kresoja-Rakic and Santoro, 2019; Yu *et al*., 2021), which is linked to the generation of RNA in the nucleus after zygotic gene activation. These potential changes in the relative and absolute profiles of RNAs and related nucleolar proteins during embryogenesis may alter the contribution of negatively charged molecules to chromatin remodeling.

### Cytoplasmic RNAs facilitate chromatin remodeling for rapid assembly of male pronucleus

In yeast, experimental increases in nuclear RNAs, achieved by inhibiting nuclear export, coincide with an increase in nuclear size (Neumann and Nurse, 2007; Kume *et al*., 2017). Additionally, in a cell-free reconstruction system using *X. laevis* egg extract, RNA degradation before initiating sperm chromatin incubation in the egg extract inhibited the assembly of nuclear envelope proteins around the chromatin (Aze *et al*., 2017). However, it remains unclear how RNAs are involved in nuclear assembly and size control. Based on the results of this study, we propose that the charge state in the egg cytoplasm, with abundant negatively charged RNAs, enhances two electrical interactions that facilitate the rapid assembly of the pronucleus from highly condensed sperm chromatin (Fig. 6G). First, the negative charge of RNAs attracts positively charged molecules, promoting the release of nuclear proteins, including protamines, from sperm chromatin. Second, the negative charge of RNAs repels negatively charged DNA or chromatin, inducing a dispersed distribution of chromatin. This chromatin remodeling by RNAs may assist histone chaperones in incorporating histones into chromatin, as excessive decompaction of chromatin by the addition of extra RNAs or negatively charged heparin increased the incorporation of histones into chromatin (Fig. 6A–C). In contrast, in the presence of RNase A in the extract, sperm nuclear proteins rarely dissociated, resulting in fewer histones being incorporated into the chromatin in the reconstructed nuclei (Fig. 6A– C). This chromatin state, with fewer nucleosomes, may explain the reported lack of accumulation of nuclear envelope proteins, such as ELYS (Aze *et al*., 2017), due to the less effective interaction between nuclear envelope proteins and the nucleosomes of the chromatin (Zierhut *et al*., 2014). Before the nuclear membrane covers the sperm chromatin, negatively charged cytoplasmic RNAs can access the negatively charged chromatin in the sperm. After the nuclear membrane forms around the sperm chromatin within 10 min (Ulbert *et al*., 2006; Franz *et al*., 2007), the remodeled chromatin, which remains in a dispersed state but with a compacted volume, generates a relatively large repulsive force to push the nuclear membrane from inside the nucleus (Hara, 2023), resulting in rapid nuclear size growth. The dispersed state of chromatin inside the nucleus, which can be experimentally manipulated by changing the epigenetic status of the chromatin, can alter the growth rate of the nucleus and its stiffness (Chan *et al*., 2017; Shimamoto *et al*., 2017; Schibler *et al*., 2023). Once the nuclear membrane with inserted nuclear pore complexes is established around the sperm chromatin, cytoplasmic RNAs—being long polymers larger than the size of the nuclear pores—and cytoplasmic negatively charged proteins no longer diffuse into the nucleus through the nuclear pores. In contrast, small ions can pass through the nuclear pores. Given the reduced permeability of RNAs through nuclear pores, the chromatin condensation state, remodeled by cytoplasmic RNAs, should be set until the nuclear membrane is fully established around the chromatin. The timing between the proposed chromatin remodeling via cytoplasmic RNAs and histone chaperone functions is expected to be coordinated for the generation of functional nuclear structures. Sperm chromatin pretreated with native RNAs or heparin, even at lower concentrations, failed to reconstruct the normal shape of the nucleus and delayed nuclear assembly (Fig. S5C). Therefore, immediately after fertilization, sperm chromatin may interact with the cytoplasmic RNAs and chaperone proteins without a temporal gap.

This study indicates that negatively charged RNAs regulate sperm chromatin remodeling in *X. laevis* fertilized eggs. The chromatin remodeled by the negative charges of RNAs is involved not only in rapid pronuclear assembly but also in other putative chromatin functions, such as transcription and the assembly of mitotic condensed chromosomes in later stages of embryonic development. The contribution of negative charges can be adjusted depending on the molecular environment of the cell. The establishment of a nuclear membrane around the chromatin reduces the accessibility of charged molecules to the chromatin through nuclear pores. Furthermore, the environment can be altered by changing the cytoplasmic composition of molecules through the acute translation of charged proteins and the local condensation of charged molecules within the cell. In eggs, the situation in which numerous RNAs surround the highly condensed sperm chromatin after fertilization is the most effective for chromatin remodeling during the short period of interphase following fertilization. Compared to general giant eggs, in smaller somatic cells, the smaller amount of RNAs per chromatin is expected to specialize in local functions, such as forming RNA-based molecular condensates like nucleoli in the cells, rather than in the drastic remodeling of the entire chromatin structure.

## Materials and methods

### Cell-free reconstruction of nucleus in *Xenopus* egg extract

Crude cytostatic factor (CSF) metaphase-arrested extracts from unfertilized *X. laevis* eggs and demembranated sperm from *X. laevis* testes were prepared as previously described (Iwabuchi *et al*., 2000; Ohsumi *et al*., 2006). Crude CSF extract was supplemented with 100 µg/ml cycloheximide (Sigma, C7698), 0.6 mM CaCl₂, 1 µM tetramethyl-rhodamine-dUTP (TMR-dUTP; Roche, 11534378910; only when visualizing replicated DNA), and 2 µM GFP–NLS recombinant proteins (only when visualizing imported proteins into the nucleus) to reconstruct the interphase nuclei.

Subsequently, demembranated sperm (∼300/µl) was mixed with the extract and incubated for the indicated duration at 22°C. The growth rate and morphology of the reconstructed nuclei differed between individual extracts due to the quality of the extract (Hara and Merten, 2015). To minimize the differences resulting from extract quality, extracts exhibiting the assembly of nuclei with aberrant morphology or those with diameters of less than 35 µm after 120 min of incubation were excluded from further experiments.

To evaluate the effects of RNA or ionic solutions on the reconstruction of the interphase nucleus in a cell-free system, the extract was supplemented with the following solutions immediately after the initiation of sperm chromatin incubation in the extract or 60 min after mixing the sperm chromatin with the extract. For conditions involving preincubation with solutions, the solutions were added to the extract and incubated for 15 min at 22°C before mixing with the sperm chromatin. For RNA degradation in the extract, RNase A (Nacalai Tesque, 30100-31) or the control buffer for RNase A (15 mM NaCl, 50%(v/v) glycerol, 10 mM Tris(3-hydroxypropyltriazolylmethyl)amine–HCl, pH 8.0) was used. To inhibit RNase A activity, RNase inhibitor (RNasIN; Promega, N2111; 1.3 U/µl) was supplemented immediately after adding 5 or 20 µg/ml RNase A into the extract containing sperm chromatin. For RNA solutions, native RNAs, purified from the *X. laevis* egg extract as described below, RNase-treated RNAs purified from egg extract treated with 100 µg/ml RNase A after 120 min of incubation, and yeast tRNAs (Thermo Scientific, AM7119) were used. Heparin sodium salt (Nacalai Tesque, 17513-96), protamine sulfate from salmon (Nacalai Tesque, 29318-41), and MgCl₂ were used as ionic solutions. To inhibit DNA replication, aphidicolin (Wako Chemicals, 011-09811; 100 µg/ml) was added immediately after mixing sperm chromatin with the extract.

### Observation of reconstructed nuclei

For measurement of the nuclear size, the extract containing the reconstructed nuclei was mixed with 10 µg/ml Hoechst 33342 (Invitrogen, H1399), 20 µg/ml 3,3’-dihexyloxacarbocyanine iodide (DiOC₆(3), Sigma, 318426), 8%(v/v) formaldehyde, and 30%(v/v) glycerol in extraction buffer (EB; 100 mM KCl, 5 mM MgCl₂, 20 mM 4-(2-hydroxyethyl)-1-piperazineethanesulfonic acid [HEPES]–KOH, pH 7.5) at a 1:1 volume ratio on the glass slide. After placing a coverslip on the extract mixture, the glass slide samples were observed using an Eclipse Ti-E (Nikon) wide-field microscope equipped with a 40× objective lens and an sCMOS camera Zyla4.2 (Andor) motorized by Micromanager. Immunostaining of the reconstructed nuclei was performed as previously described (Hara and Merten, 2015) using anti-histone H3 antibody (MBL, MABI0301, ×1,000 dilution), anti-histone H2A antibody (Abcam, ab1970, ×200 dilution), supernatant of hybridoma against nucleolin (Developmental Studies Hybridoma Bank, clone:p7-1A4 and b6-6E7, ×25 dilution), anti-fibrillarin antibody (Abcam, ab5821, ×1,000 dilution), anti-α-tubulin antibody (DM1A clone; Thermo Fisher Scientific, MS581P0, ×1,000 dilution), Alexa 633 anti-mouse IgG antibody (Thermo Fisher Scientific, A121050, ×1,000 dilution), Alexa 633 anti-rabbit IgG antibody (Thermo Fisher Scientific, A21070, ×1,000 dilution), 10 µg/ml Hoechst 33342, and 10 µg/ml DiOC₆(3). For tubulin immunostaining, extracts containing the reconstructed nuclei were fixed and stained under different sedimentation conditions to immunostain histones and nucleolar proteins, as described previously (Hara and Merten, 2015). Immunostained nuclei were observed using a FV1200 (Olympus) confocal laser microscope equipped with a 60× objective lens and an Eclipse Ti-E wide-field microscope equipped with a 20× objective lens to visualize histones or nucleolar proteins and tubulin, respectively. For RNA staining, the extract containing the reconstructed nuclei, after 60 min of incubation or isolated nuclei from embryos, was mixed with EB containing 10 µg/ml Hoechst 33342, 8%(v/v) formaldehyde, 30%(v/v) glycerol, and RNA staining dye by a 1:1 volume ratio on the slide. Pyronin Y (Cayman Chemical, 14488; 330 µM), SYTO RNA Select Green (Thermo Scientific, S32703; 0.5 µM), or Strand Bright (AAT Bioquest, 17610; 1:1000 dilution) was used.

### Isolation of RNAs from samples

Total RNA was isolated from egg extracts or embryos using TRI reagent (Merck, 93289) according to the manufacturer’s instructions. Briefly, egg extracts containing reconstructed nuclei were mixed with TRI reagent. For RNA isolation from embryos, unfertilized eggs from female *X. laevis* were inseminated with homogenized testes isolated from male *X. laevis*. After removing the egg jelly, the fertilized eggs were incubated until the indicated developmental stages, as described previously (Shimogama *et al*., 2022). Following incubation, 20 embryos were mixed with TRI reagent by pipetting. The mixture was frozen at –80°C and supplemented with chloroform. After centrifugation, the aqueous phase was transferred to a fresh microtube containing 70% isopropanol. Following another centrifugation, the RNA pellet was dissolved in nuclease-free H₂O and adjusted to the desired concentration for further experiments. To confirm RNA profiles, the isolated RNAs were subjected to agarose gel (1–5% (v/v)) electrophoresis.

### Isolation of nuclei from embryos

The fertilized eggs were incubated until the gastrula embryonic stages for 17 h at 16°C in 0.2× MMR (1× MMR: 0.25 mM HEPES-KOH, 0.1 mM ethylenediamine-N,N,N’,N’-tetraacetic acid, 5 mM NaCl, 2 mM KCl, 1 mM MgCl₂, 2 mM CaCl₂, pH 7.8). The embryos at the desired stages were incubated in 0.2× MMR containing 100 µg/ml cycloheximide for 60 min at 22°C to block the cell cycle at interphase. After washing the embryos twice with ice-cold EB and ice-cold EB containing 250 mM sucrose, they were homogenized using a conventional P200 pipette. The mixture was placed in ice-cold EB containing 1.3 M sucrose in a microtube and centrifuged using a swing rotor at 2,000 × *g* for 3 min at 4°C. The black pellet on the 1.3 M sucrose cushion was transferred to a new microtube and mixed with nearly the same volume of ice-cold EB. After centrifugation of the mixture at 500 × *g* for 3 min at 4°C, the pellet containing the isolated nuclei was dissolved in EB containing 10%(v/v) glycerol.

### Incubation of sperm chromatin and isolated nuclei with the solution

Demembranated sperm or nuclei isolated from developed embryos were mixed with RNAs or ionic solutions. After incubating the mixture for the indicated duration at 22°C, the mixture was fixed and stained with EB containing 10 µg/ml Hoechst 33342, 8%(v/v) formaldehyde, and 30%(v/v) glycerol by a 1:1 volume ratio on the slide glass for microscopic observation. To examine the reversibility of the chromatin compaction state, the sperm chromatin was first incubated with the ionic or RNA solution for 10 min, after which 100 µg/ml RNase A or 1 mg/ml protamine was added to the preincubated sample, and the mixtures were incubated for an additional 10 min. For detection of proteins extracted from the chromatin, the incubated mixture of sperm chromatin was centrifuged at 20,000 × *g* for 10 min at 25°C. The supernatant was transferred to a new microtube and mixed with SDS sample buffer (Wako Chemicals, 193-11032) containing 10% 2-mercaptoethanol. After incubation at 100°C for 15 min, the mixture was subjected to SDS-PAGE. The SDS-PAGE gel was stained with CBB Stain One (Wako Chemicals, 11642-31).

### Quantification and statistical analysis of nuclear parameters

To measure the nuclear size, the cross-sectional area of the reconstructed nucleus was assessed for the fixed sample. At least 20 nuclei from at least two individual extracts were analyzed for each experimental condition. To minimize the effects of variations in extract preparation, only data showing reconstructed nuclei with a diameter greater than 35 µm in the control condition (without any additional solutions) after 120 min of incubation were used for further analysis. For normalization of each extract, the measured nuclear cross-sectional areas at a particular incubation time and condition were divided by the mean values obtained after 60 or 120 min of incubation under control conditions using the same extract preparation. The growth rate was calculated from the slope of the regression line of the mean nuclear cross-sectional area plot for each extract preparation after 40–120 min of incubation. The mean values for each condition were calculated from the mean nuclear parameters and standard deviations from the individual experiments, and these were illustrated in the graphs.

For quantification of the signal intensity of immunostained proteins within the nucleus, the immunostained samples were prepared under the same conditions as the immunostaining, and the images were acquired using the same microscopy parameters for all samples being compared. From these images, the mean intensities of the immunostained proteins and DNA stained with Hoechst 33342 per pixel inside the nuclear region were measured using ImageJ with the ellipsoid function and subtracted from the background. Additionally, the intensities inside the nucleus were calculated by multiplying the mean intensity by the measured cross-sectional area. The relative intensity of the immunostained proteins was calculated by dividing the total intensity of the immunostained proteins by that of the DNA in each nucleus. To normalize the variation among the extract preparations, the calculated relative intensities were divided by the mean intensity values of the nuclei under control conditions.

To quantify the imported GFP-NLS proteins within the nucleus and incorporated TMR-dUTP within the chromatin, the mean intensity of GFP-NLS or TMR-dUTP within the nuclear region was measured using ImageJ with the ellipsoid function and subtracted from the background. The intensities inside the nucleus were also calculated by multiplying the mean intensity by the measured cross-sectional area. To analyze the speed of DNA replication, the relative intensity of TMR-dUTP was calculated by dividing the intensity of TMR-dUTP at each time point of incubation by the mean value of the total intensity of TMR-dUTP from the control sample after 120 min of incubation under control conditions for each extract preparation.

To calculate the entropy of chromatin within the nucleus, images of the reconstructed nuclei stained with Hoechst 33342 were acquired under the same staining and exposure conditions for all samples being compared. The images were then converted to 8-bit images by setting the lowest and highest intensity values within the nucleus to 0 and 255, respectively. From the images, the intensity profiles of Hoechst 33342-positive intensity within the observed cross-section of the reconstructed nuclei after 120 min of incubation were obtained using the “plot profile” function in ImageJ. Using the intensity profile, the probability distribution of the intensity was calculated using the formula: Entropy = Σ P(x) ln (P(x)), where P(x) represents the probability of each intensity.

Significant differences among the samples were determined by non-parametric Wilcoxon signed-rank tests using the R software. To analyze the differences among histone relative intensities with a large sample size, Student’s t-test was performed using the function available in Excel software.

## Acknowledgements

We are grateful to Misako Ohama, Shuichi Nakano, and Tomonari Kato in our laboratory for preparing the experimental setup for this study. This work was supported by JSPS KAKENHI, Grant Numbers JP21H00256, JP21K19268, and JP23H04287.

