## supplemental figures for "Rapid pronucleus assembly using cytoplasmic RNAs in fertilized eggs of *Xenopus laevis*"

Mizuki Ikeda, Yuto Tanaka, Tatsuya Shohoji, Yuki Hara.
Evolutionary Cell Biology Laboratory, Faculty of Science, Yamaguchi University
Yoshida 1677-1, Yamaguchi city, 753-8512, Japan.

### SUPPLEMENTAL FIGURES

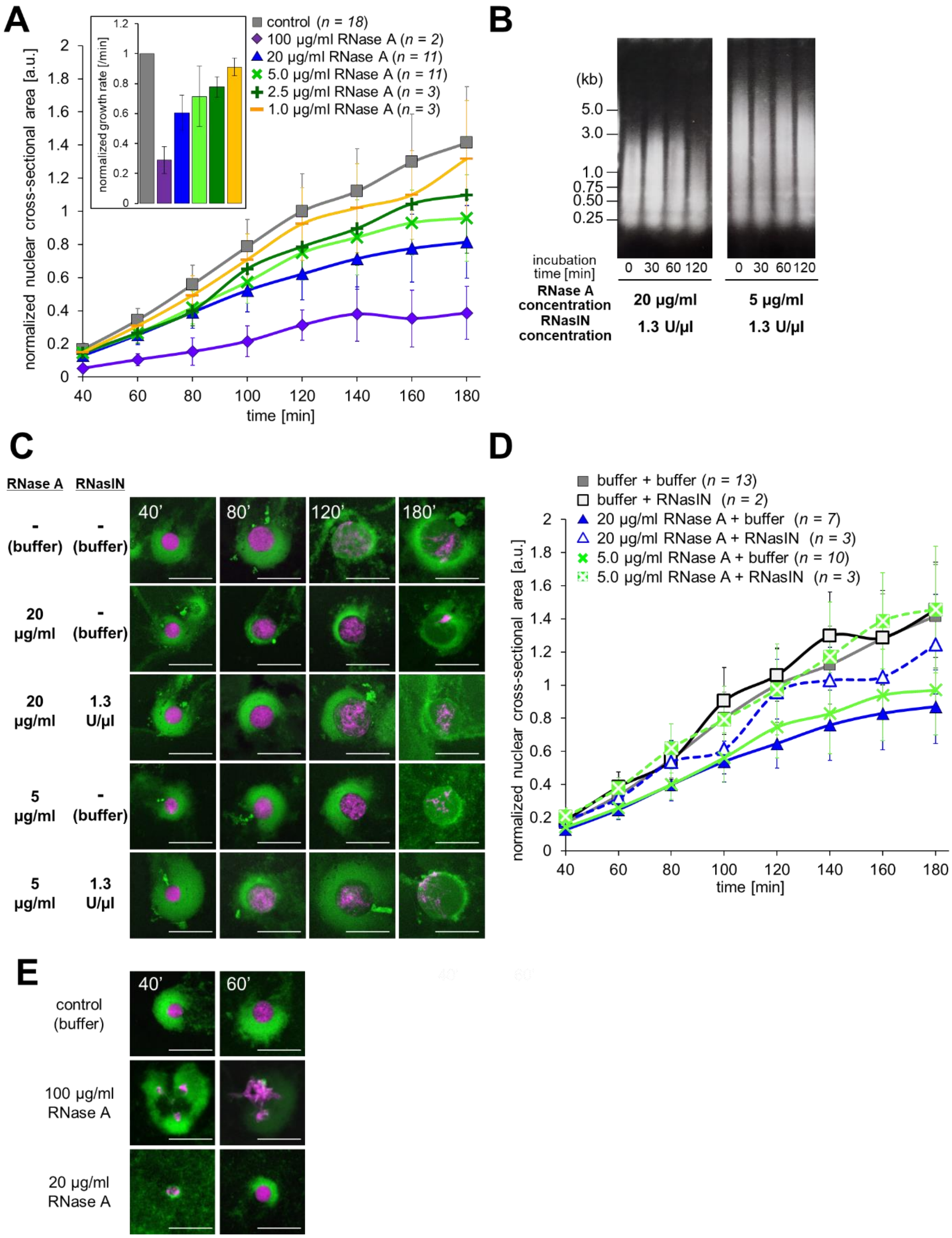


**Figure S1.** (**A**) Dynamics of the mean normalized cross-sectional areas of the nuclei reconstructed from *X. laevis* egg extract with each concentration of RNase A. Mean nuclear cross-sectional areas at each incubation period was normalized by the mean values in the control condition after 120 min of incubation. Average of the normalized mean nuclear cross-sectional areas (±SD) are connected using a line in each condition with different RNase A concentration. The inset shows the average of normalized growth rates of nuclear cross-sectional area (±SD) in each RNase A concentration. Numbers (*n*) of experiments using each extract preparation for calculating the growth rate were shown. The graph displaying the absolute values were shown in Fig. 1D. (**B**) The purified total RNAs from *X. laevis* egg extract supplemented with RNase A and RNase inhibitor (RNasIN) of the indicated concentration and incubated for the indicated duration were analyzed by agarose gel electrophoresis. The nucleotide sizes were shown based on the known DNA ladder on the left. (**C**) Representative images of the nuclei reconstructed in the *X. laevis* egg extract in the presence of RNase A and RNasIN of the indicated concentration. The samples were stained with Hoechst 33342 (for visualizing DNA; magenta) and $DiOC_6(3)$ (for visualizing membrane; green). Scale bars, 50 µm. (**D**) Dynamics of the normalized mean cross-sectional areas of the nuclei reconstructed from *X. laevis* egg extract with each concentration of RNase A and RNasIN. Averages of normalized mean cross-sectional areas (±SD) at each incubation period are connected using a line in each condition. (**E**) Representative images of the nuclei reconstructed in the *X. laevis* egg extract with pre-incubation with RNase A at the indicated concentration for 15 min before mixing the extract with sperm chromatin. The samples were stained with Hoechst 33342 (for visualizing DNA; magenta) and $DiOC_6(3)$ (for visualizing membrane; green).

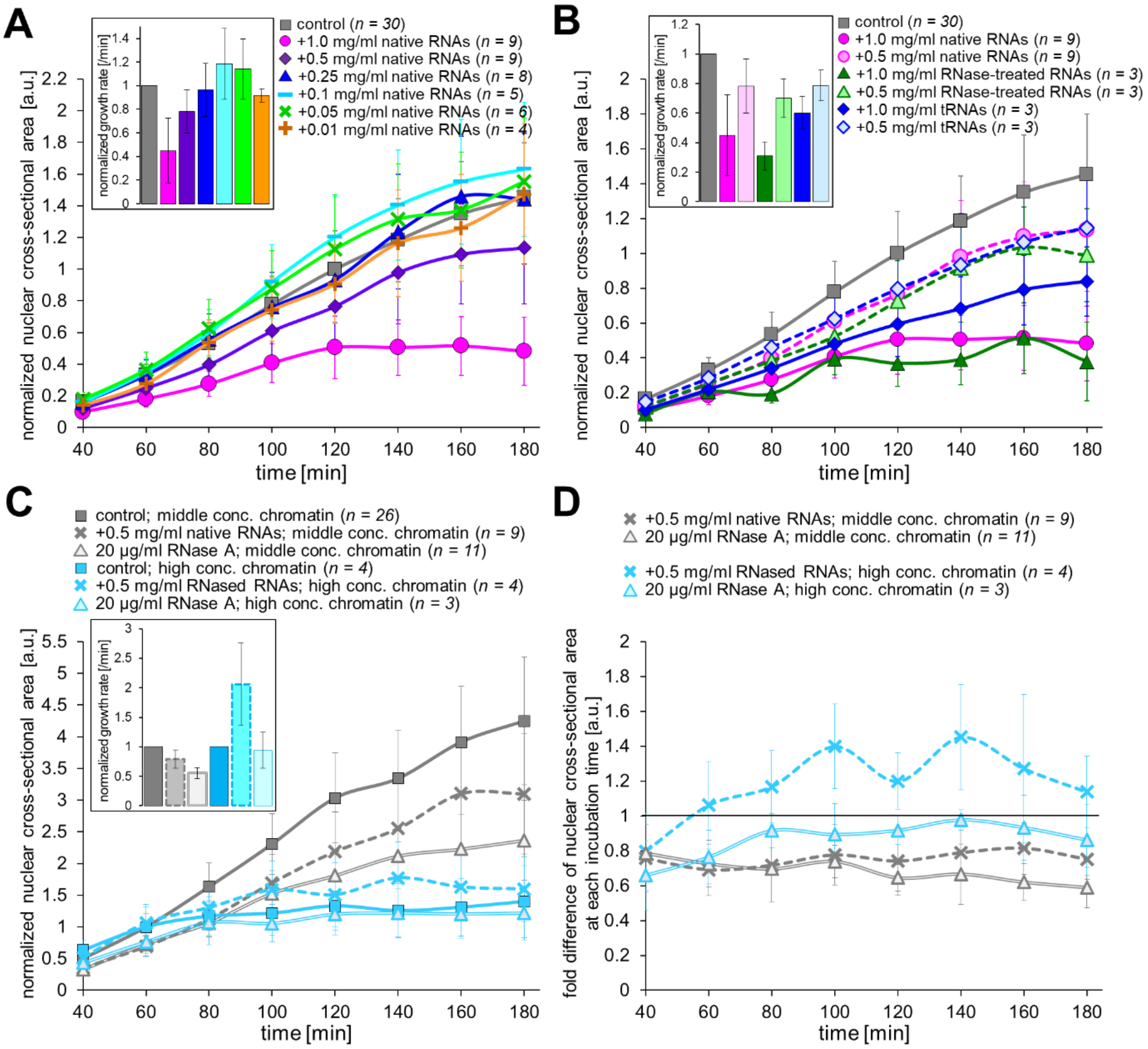


**Figure S2.** (**A-C**) Dynamics of the mean normalized cross-sectional areas of the nuclei reconstructed from *X. laevis* egg extract with adding native RNAs at various concentrations (**A**), native RNAs, RNased RNAs, or tRNAs at various concentrations (**B**), or plus 0.5 mg/ml native RNAs and 20 µg/ml RNase A with various sperm concentration (**C**). Mean nuclear cross-sectional areas at each incubation period was normalized by the mean values in the control condition (without extra RNA or RNase A) after 120 min (**A, B**) or 60 min (**C**) of incubation. In the condition of different sperm concentration, the normalization was performed in each condition of sperm concentration. Average of the normalized mean nuclear cross-sectional areas (±SD) are connected using a line in each condition. The inset shows the average of normalized growth rates of nuclear cross-sectional area (±SD) in each condition. Numbers (*n*) of experiments using each extract preparation for calculating the growth rate were shown. The graphs displaying the absolute values were shown in Fig. 2B, 2D, 2E. (**D**) Dynamics of the fold difference of the mean cross-sectional areas in each condition with addition of extra RNAs or RNase A. Error bar represents the mean relative SD value by dividing the measured SD by the mean nuclear cross-sectional areas in the control condition at each incubation period.

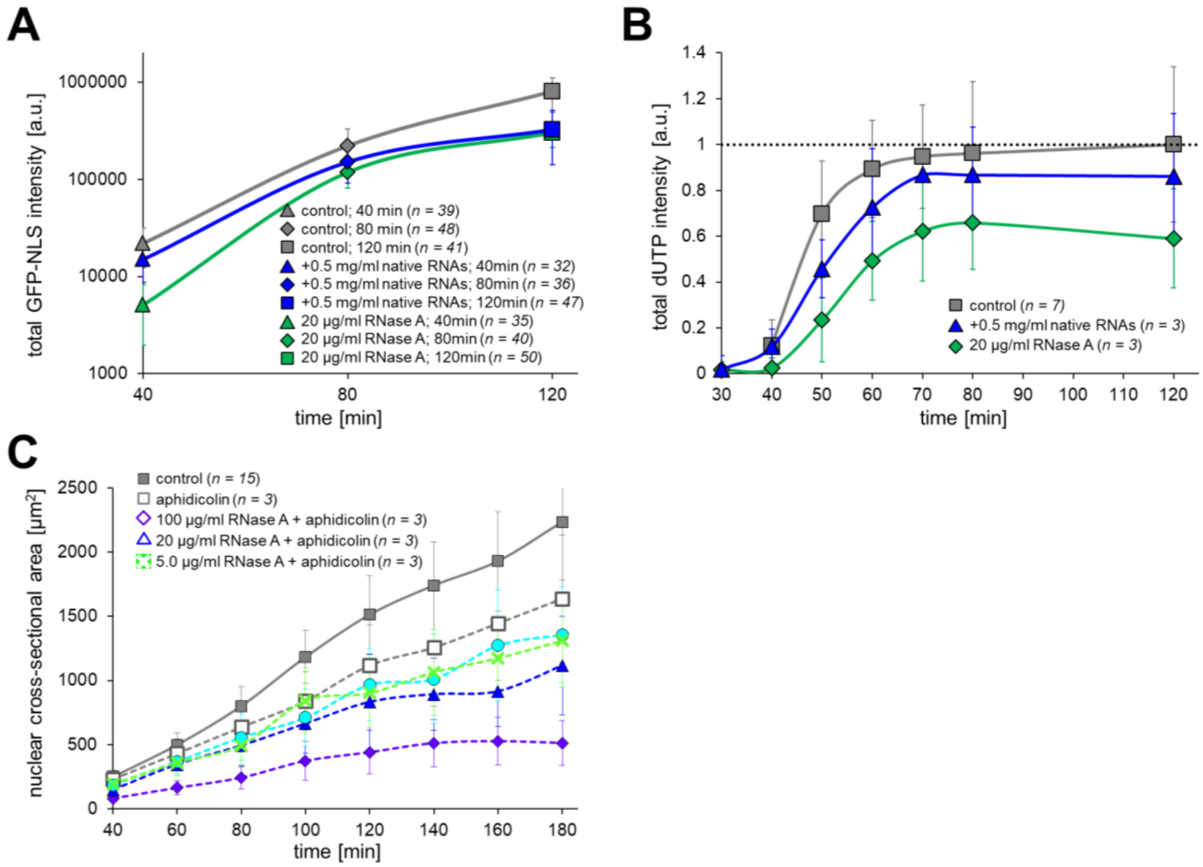


**Figure S3.** (**A**) Dynamics of the total intensity of GFP-NLS in the cross-sectional areas of the nuclei reconstructed from *X. laevis* egg extract with adding plus 0.5 mg/ml native RNAs or 20 µg/ml RNase A. Average of the calculated total intensities (± SD) are connected using a line in each condition. Numbers (*n*) of observed samples for measuring GFP-NLS intensity were shown. (**B**) Dynamics of the normalized total intensity (± SD) of TMR-dUTP within the cross-sectional areas of the nuclei reconstructed from *X. laevis* egg extract with adding plus 0.5 mg/ml native RNAs or 20 µg/ml RNase A. The calculated total intensity was normalized by that in the control condition after 120 min of incubation. Numbers (*n*) of experiments using each extract preparation for measuring the TMR-dUTP intensity were shown. (**C**) Dynamics of the mean cross-sectional areas (± SD) of the nuclei reconstructed from *X. laevis* egg extract with adding plus 0.5 mg/ml native RNAs or 20 µg/ml RNase A and DNA replication inhibitor, aphidicolin.

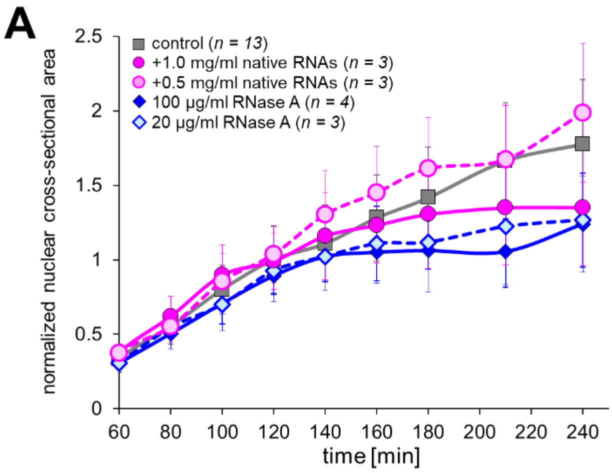


**Figure S4.** (**A**) Dynamics of the mean normalized cross-sectional areas of the nuclei reconstructed from *X. laevis* egg extract with adding native RNAs or RNase A at various concentrations after incubation of egg extract with sperm concentration for 60 min. Mean nuclear cross-sectional areas at each incubation period was normalized by the mean values in the control condition (without extra RNA or RNase A) after 120 min. Average of the normalized mean nuclear cross-sectional areas (±SD) are connected using a line in each condition. Numbers (*n*) of experiments using each extract preparation for measuring the nuclear cross-sectional area were shown. The graph displaying the absolute values were shown in Fig. 4B.

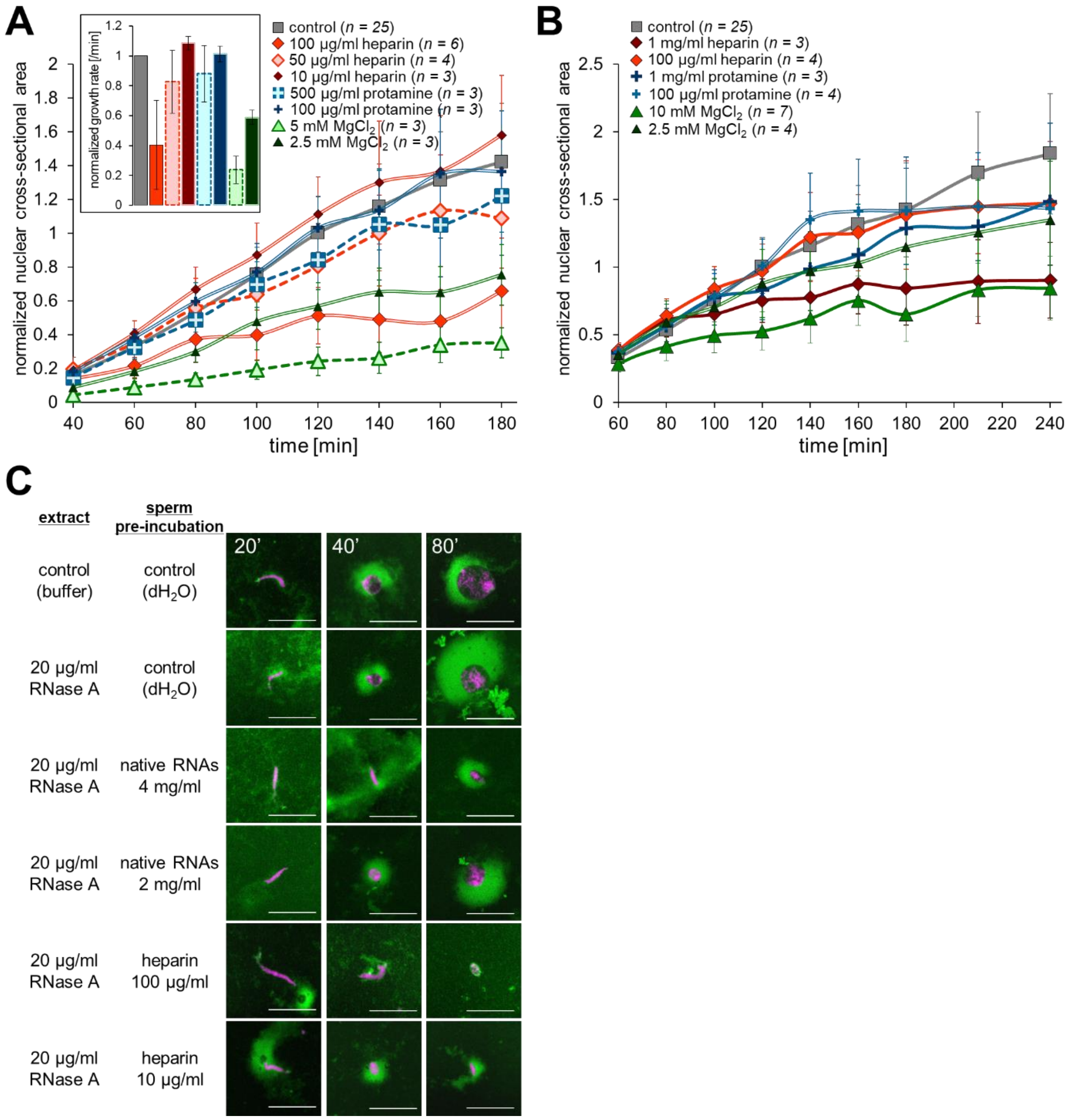


**Figure S5.** (**A**, **B**) Dynamics of the mean normalized cross-sectional areas of the nuclei reconstructed from *X. laevis* egg extract with adding ionic solutions at various concentrations simultaneously with sperm chromatin (**A**) or after 60 min of incubation with sperm chromatin (**B**). Mean nuclear cross-sectional areas at each incubation period was normalized by the mean values in the control condition (without ionic solutions) after 120 min of incubation. Average of the normalized mean nuclear cross-sectional areas (±SD) are connected using a line in each condition. The inset shows the average of normalized growth rates of nuclear cross-sectional area (±SD)

in each condition. Numbers (*n*) of experiments using each extract preparation for calculating the growth rate were shown. The graphs displaying the absolute values were shown in Fig. 5E, 5F. (**C**) Representative images of the chromatin after incubation of sperm chromatin, which is preincubated with solution containing with native RNAs or heparin at different concentrations, with *X. laevis* egg extract supplemented with 20 µg/ml RNase A. The extract containing the sperm chromatin was stained with Hoechst 33342 (magenta) and $DiOC_6(3)$ (for visualizing membrane; green) at the indicated incubation period. Scale bars, 50 µm.

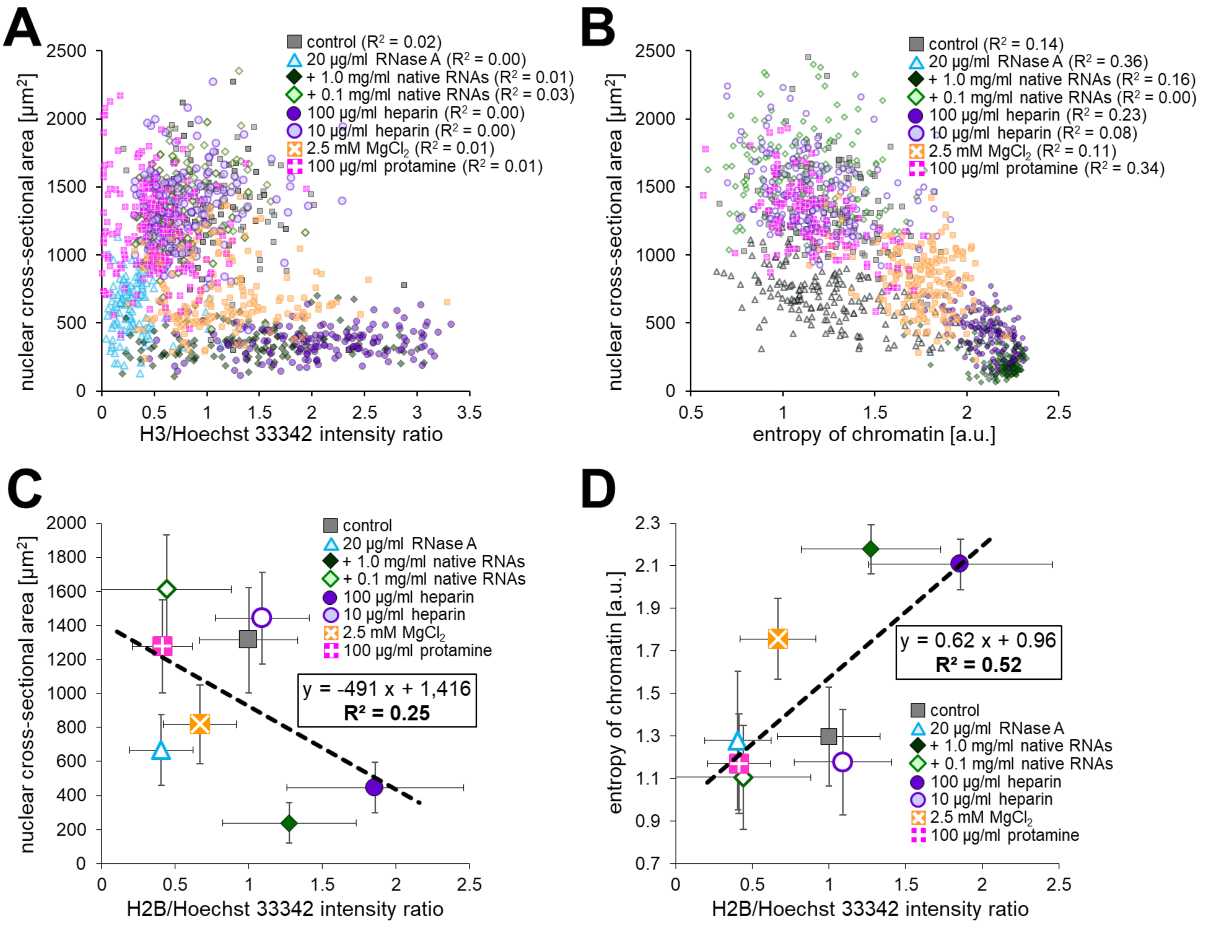


**Figure S6.** (**A**) Scatter plots of the measured individual nuclear cross-sectional areas in each condition of the indicated supplements with the intensity ratio of histone H3/Hoechst 33342. (**B**) Scatter plots of the calculated individual value of the chromatin entropy in each condition of the indicated supplementation with the relative intensity ratio of histone H3/Hoechst 33342. The determination of the correlation ($R^2$) from the data in each condition was shown. (**C**) Scatter plots of the measured mean nuclear cross-sectional areas in each condition of the indicated supplements with the intensity ratio of histone H2B/Hoechst 33342. (**D**) Scatter plots of the calculated value of the chromatin entropy in each condition of the indicated supplementation with the relative intensity ratio of histone H2B/Hoechst 33342. Dataset from the nuclei reconstructed in the egg extract (colored symbols) was fitted using a linear regression with showing the equation and the value of determination of the correlation ($R^2$).
